# Genome-wide misexpression associated with hybrid sterility in *Mimulus* (monkeyflower)

**DOI:** 10.1101/750687

**Authors:** Rachel E. Kerwin, Andrea L. Sweigart

## Abstract

Divergence in gene expression regulation is common between closely related species and may give rise to incompatibilities in their hybrid progeny. In this study, we investigated the relationship between regulatory evolution within species and reproductive isolation between species. We focused on a well-studied case of hybrid sterility between *Mimulus guttatus* and *M. nasutus*, two closely related yellow monkeyflower species, that is caused by two epistatic loci, *hybrid male sterility 1* (*hms1*) and *hybrid male sterility 2* (*hms2*). We quantified and compared global transcript abundance across male and female reproductive tissues (*i.e.* stamens and carpels) of *M. guttatus* and *M. nasutus*, as well as sterile and fertile progeny from an advanced *M. nasutus*-*M. guttatus* introgression line that carries the *hms1-hms2* incompatibility. We observed substantial variation in transcript abundance between *M. guttatus* and *M. nasutus*, including distinct but overlapping patterns of tissue-biased expression, providing evidence for regulatory divergence between these species. Furthermore, we found pervasive genome-wide misexpression exclusively associated with hybrid sterility – only observed in the affected tissues (*i.e.* stamens) of sterile introgression hybrids. Examining patterns of allele-specific expression in sterile and fertile hybrids, we found evidence of *cis-* and *trans-* regulatory divergence, as well as *cis-trans* compensatory evolution (likely to be driven by stabilizing selection). However, regulatory divergence does not appear to cause misexpression in sterile hybrids, which instead likely manifests as a downstream consequence of sterility itself.

## INTRODUCTION

Closely related species often show considerable regulatory divergence – that is, they have accumulated differences in the *cis*-acting DNA sequences or *trans*-acting factors that regulate gene expression (Tautz, 2000; Wray *et al.*, 2003). As with any epistatic loci, divergence in interacting regulatory elements might lead to incompatibilities in the hybrid progeny of interspecific crosses (Dobzhansky, 1937; Muller, 1942; Mack & Nachman, 2017). These hybrid incompatibilities might arise due to independent substitutions in distinct lineages, with genetic drift or selection increasing the frequency of a derived allele at a *cis*-acting locus in one species and a *trans*-acting partner locus in the other. In the classic Dobzhansky-Muller Model, these derived alleles are neutral or favored on their own, but cause aberrant gene expression when combined in hybrids. Alternatively, hybrid incompatibilities might arise because of coevolution between *cis*- and *trans*-elements within a single lineage. Stabilizing selection to maintain optimal levels of transcription, which favors *cis*- and *trans*-regulatory variants that compensate for each other, appears to be an important force shaping gene expression evolution (Gilad *et al.*, 2006; Tirosh *et al.*, 2009; Goncalves *et al.*, 2012; Coolon *et al.*, 2014; Mack *et al.*, 2016). Thus, even when transcript abundance for a particular gene does not differ between species, the underlying regulatory components controlling its expression might have diverged (Tautz, 2000; True & Haag, 2001; Wray *et al.*, 2003). This process of compensatory evolution in gene regulatory networks, which is likely to affect different sets of genes in diverging lineages, has been proposed as a major source of hybrid incompatibilities between species (Landry *et al.*, 2005; Takahasi *et al.*, 2011).

Despite the clear importance of changes in gene expression for phenotypic evolution (Wittkopp, 2013), empirical support for regulatory divergence as a general driver of hybrid incompatibilities is mixed. While many studies have uncovered pervasive gene misexpression in sterile hybrids (Michalak & Noor, 2003; Ranz *et al.*, 2004; Haerty & Singh, 2006; Malone *et al.*, 2007; Coolon *et al.*, 2014; Brill *et al.*, 2016; Mack *et al.*, 2016), others have found no association between patterns of gene expression and hybrid dysfunction (Barbash & Lorigan, 2007; Wei *et al.*, 2014; Guerrero *et al.*, 2016). Even when sterile or inviable hybrids do show misexpression, it is usually difficult to determine which is cause and which is effect. Because hybrid dysfunction often involves gross defects in affected tissues (e.g., testes in male sterile hybrids), global gene misregulation might occur as a downstream consequence of abnormal or missing cell types. For example, although misexpression increases dramatically in hybrids between *Drosophila* species pairs with longer divergence times (Coolon *et al.*, 2014), so does the severity of hybrid dysfunction.

A promising approach to disentangle the causes of hybrid incompatibilities from their downstream effects is to examine interspecific gene expression variation associated with particular genomic regions. Although most studies of regulatory divergence compare gene expression profiles between parental species and F1 hybrids, a handful have used introgression lines (Lemos *et al.*, 2008; Meiklejohn *et al.*, 2014; Guerrero *et al.*, 2016) or recombinant mapping populations (Turner *et al.*, 2014), which can facilitate investigations into whether gene regulation and hybrid dysfunction have a shared genetic basis. With the introgression approach, it is possible to examine the regulatory effects of small genomic regions from one species on the genetic background of another species, and vice versa. Additionally, comparing the regulatory effects of introgressions with and without hybrid incompatibility phenotypes can address the generality of misregulation as a cause of hybrid dysfunction (Guerrero *et al.*, 2016).

To investigate the link between regulatory divergence and hybrid dysfunction, we exploited a well-studied hybrid incompatibility system between two closely related species of monkeyflower (*Mimulus*). Previously, we discovered severe male sterility and partial female sterility in hybrids between *Mimulus guttatus* (IM62 inbred line) and *M. nasutus* (SF5 inbred line) (Sweigart *et al.*, 2006) and fine-mapped the effects to two small nuclear genomic regions of ~60 kb each on chromosomes 6 and 13 (Sweigart & Flagel, 2015). Hybrids that carry at least one incompatible *M. guttatus* IM62 allele at *hybrid male sterility 1* (*hms1*) on chromosome 6 in combination with two incompatible *M. nasutus* SF5 alleles at *hybrid male sterility 2* (*hms2*) on chromosome 13 display nearly complete (>90%) pollen sterility, whereas other allelic combinations are highly fertile (Sweigart *et al.*, 2006; Kerwin & Sweigart, 2017). Here, we took advantage of SF5-IM62 introgression hybrids, formed through multiple rounds of selection for pollen sterility and backcrossing (as the female parent) to *M. nasutus* SF5 (Figure 1). This *r*ecurrent *s*election with *b*ackcrossing (RSB) population maintains a heterozygous IM62 introgression on chromosome 6 (that contains the *hms1* locus) against an otherwise SF5 genetic background (including at *hms2*). Each generation, RSB progeny segregate ~1:1 for sterility to fertility, depending on whether they inherit a copy of the incompatible IM62 *hms1* allele. Additionally, nearly all RSB hybrids are expected to carry a heterozygous IM62 introgression on chromosome 11 surrounding the female meiotic drive locus *D*, which is transmitted to >98% of progeny when an SF5-IM62 F1 hybrid acts as the maternal parent (Fishman & Willis, 2005). Thus, this crossing scheme produces two classes of progeny: sterile (STE) individuals that carry two heterozygous introgressions (on chromosome 6 with *hms1* and on chromosome 11 with *D*) and fertile (FER) individuals that carry a single introgression (on chromosome 11 with *D*). The result is an internally controlled genetic experiment that is ideally suited for determining whether gene misexpression is a cause or consequence of hybrid sterility.

**Figure 1.**
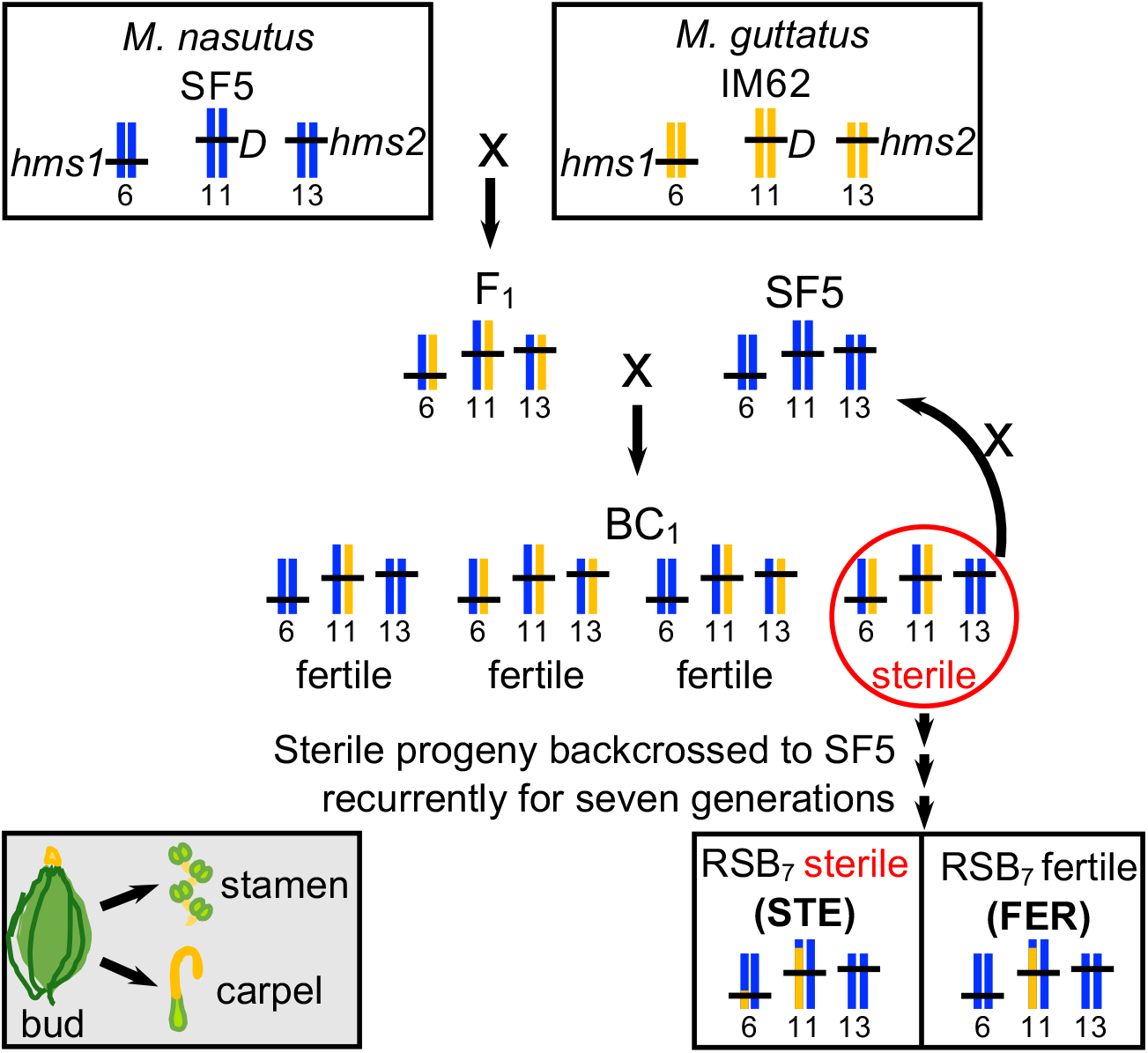
Crossing scheme to generate *r*ecurrent *s*election with *b*ackcrossing (RSB) population. First, an SF5 x IM62 F_1_ was backcrossed to SF5, yielding a first generation backcross (BC_1_) population. A pollen sterile individual from the BC_1_ population (red circle) was backcrossed to SF5, yielding a first generation introgression line (RSB_1_). The selective backcrossing was repeated for six more generations to produce an RSB_7_ population. Roughly 50% of the RSB_7_ siblings are pollen sterile because they carry an heterozygous introgression of the incompatible IM62 allele at *hms1* in an SF5 genomic background that is fixed for the incompatible allele at *hms2* while the other 50% are pollen fertile because they carry an SF5 allele at *hms1* in the same genomic background. Whole transcriptome sequencing was performed on three bioreps each of stamens and carpels (grey box) from four genotypes, *M. nasutus* SF5, *M. guttatus* IM62, RSB_7_ fertile (FER), and RSB_7_ sterile (STE) (black boxes), for a total of 24 samples representing eight tissue by genotype categories. To obtain sufficient tissue for sequence library prep, carpels or stamens dissected from four to eight young buds were pooled to form a single biorep.

For this study, we sequenced the transcriptomes of STE and FER progeny from a seventh-generation RSB population, (*i.e.* RSB_7_) alongside their parents, *M. guttatus* IM62 and *M. nasutus* SF5. Because the *hms1-hms2* incompatibility affects both male and female fertility (Sweigart *et al.*, 2006), we isolated RNA separately from developing stamen and carpel tissue. This RNAseq dataset allowed us to identify changes in gene expression due to the presence of a genomic segment that includes a known hybrid sterility allele (*i.e*., the IM62 allele at *hms1*) versus a genomic segment that does not (*i.e*., the IM62 allele at *D*). If regulatory divergence between *Mimulus* species is substantial, we would expect to see expression differences induced by both introgressions. In particular, the introgression lines might show transgressive expression – genes expressed outside of the parental range – due to heterospecific combinations of divergent *cis*- and *trans*-acting factors. If instead, transcriptional misregulation is confined to sterile hybrid samples, it might suggest modest regulatory divergence between species and large downstream effects of the *hms1-hms2* incompatibility. With this RNAseq dataset, we addressed the following specific questions. To what extent does gene expression vary between closely related *Mimulus* species? Is there evidence for tissue-biased gene expression between the stamens and carpels? What are the mechanisms of regulatory divergence between species? Is there an association between transcriptional misregulation and hybrid sterility and, if so, what is its cause? Do expression patterns narrow down the list of candidate genes for *hms1* or *hms2*? Our results provide insight into regulatory divergence between closely related species and the consequences for reproductive isolation.

## RESULTS

To examine patterns of genome-wide expression associated with the *hms1-hms2* hybrid incompatibility in *Mimulus*, we performed transcriptome sequencing (RNAseq) of stamens and carpels from *M. guttatus* IM62 and *M. nasutus* SF5, as well as from fertile (FER) and sterile (STE) siblings of an advanced (seventh generation) SF5 x IM62 introgression population called *r*ecurrent *s*election with *b*ackcrossing (RSB_7_) (Figure 1; see Methods for more details on crossing scheme). We obtained an average of 14.1 million (range: 10.9 - 16.8 million) 50-bp single-end sequencing reads from each sample (Table S1). After initial pre-processing, we aligned trimmed reads to either the *M. guttatus* v2.0 reference genome (http://phytozome.net) or a pseudoreference SF5 genome generated for this study (see Methods for details). An average of 77.4% (range: 64.5 – 88.3%) of the reads mapped uniquely, and no more than 5.3% mapped to multiple locations (Table S1). Across the 14 chromosomes, we found that 21,147 of 28,140 predicted genes were expressed (i.e. read counts per million [CPM] >1 in >= 3 samples).

To determine introgression boundaries in the STE and FER RSB_7_ siblings, we called genome-wide single nucleotide polymorphisms (SNPs) and used them to genotype the samples. As expected due to our crossing scheme (see Figure 1), STE and FER siblings differ only in the region surrounding *hms1* on chromosome 6. Here, STE individuals contain a heterozygous IM62 introgression that stretches across a 7 Mb region encoding 698 genes (508 of which are expressed in our dataset) while FER individuals are homozygous for the recurrent SF5 parent (Figure S1). Additionally, both FER and STE individuals carry a large heterozygous IM62 introgression that spans 23 Mb of chromosome 11 (90% of the physical distance; Figure S1), a region that encodes 1064 genes (800 expressed in our dataset) and harbors a female meiotic drive locus (*D*) associated with strong transmission ratio distortion in SF5 x IM62 hybrids (Fishman & Willis, 2005; Fishman & Saunders, 2008).

### Expression variation is driven by species, tissue, and fertility

To visualize global gene expression patterns across the 24 samples in our dataset, we generated a multidimensional scaling (MDS) plot (Figure 2). Replicate samples from each of the eight genotype-by-tissue groups are highly similar to one another and form clusters that are generally separated from other groups (Figure 2). Along the y-axis, samples are primarily separated by species identity, with *M. guttatus* IM62 showing a distinct pattern of expression from *M. nasutus* SF5 and siblings from the RSB_7_ introgression line (the latter carrying SF5 variation across most of their genetic backgrounds, see Figure S1). Of the 21,147 expressed genes, roughly 9% were significantly differentially expressed (−2 < log2 fold-change > 2, FDR ≤ 0.05) between IM62 and SF5 in each tissue type (Figure S2). Previous work has shown substantial nucleotide divergence between *M. guttatus* and *M. nasutus* (*d*_*S*_ between IM62 and SF5 is 4.5%, see Brandvain et al. 2014), which presumably accounts for much of this expression variation.

**Figure 2.**
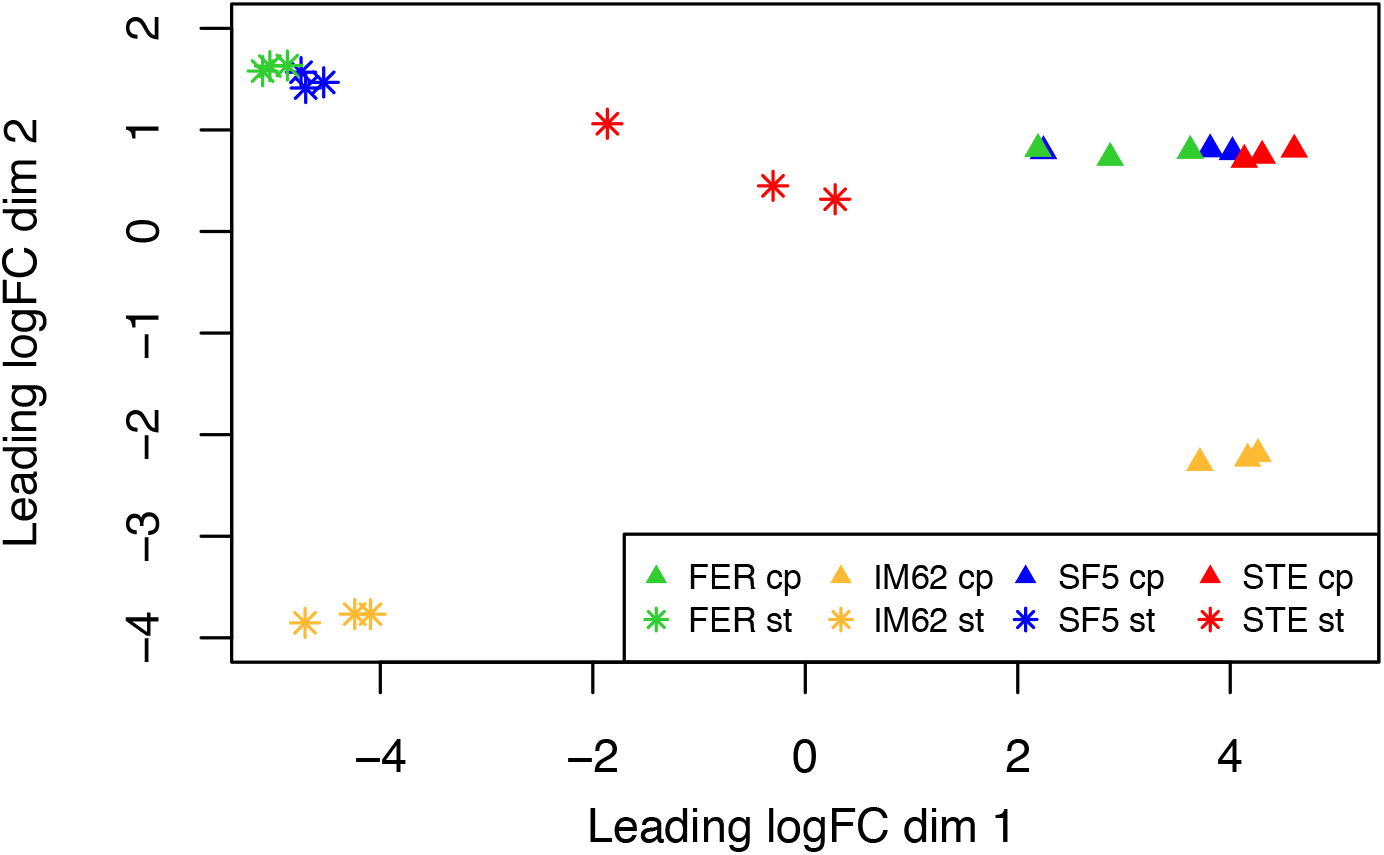
Genome-wide expression pattern across samples. Plot shows results from multidimensional scaling (MDS) analysis comparing gene expression across all 24 RNAseq samples. The colors and shapes represent the different genotype by tissue sample categories. FER cp = FER carpel, FER st = FER stamen, IM62 cp = IM62 carpel, IM62 st = IM62 stamen, SF5 cp = SF5 carpel, SF5 st = SF5 stamen, STE cp = STE carpel, STE st = STE stamen, logFC = log-fold-change

Along the x-axis of the MDS plot, expression variation is largely determined by tissue type with clear separation between carpel and stamen samples (Figure 2). To investigate these differences more thoroughly, we compared patterns of tissue-biased gene expression between IM62 *M. guttatus* and SF5 *M. nasutus* (Figure 3). For a large number of genes, tissue-biased expression is conserved between species: 3393 genes (16%) show higher/lower expression in the same tissues in both IM62 and SF5 (green points in Figure 3). On the other hand, a considerable number of genes (3214, 15.2%; blue and yellow dots in Figure 3) show tissue-biased expression in only one of the two species and a handful (66, 0.4%; purple points in Figure 3) even show opposite patterns of tissue-biased expression (Figure 3).

**Figure 3.**
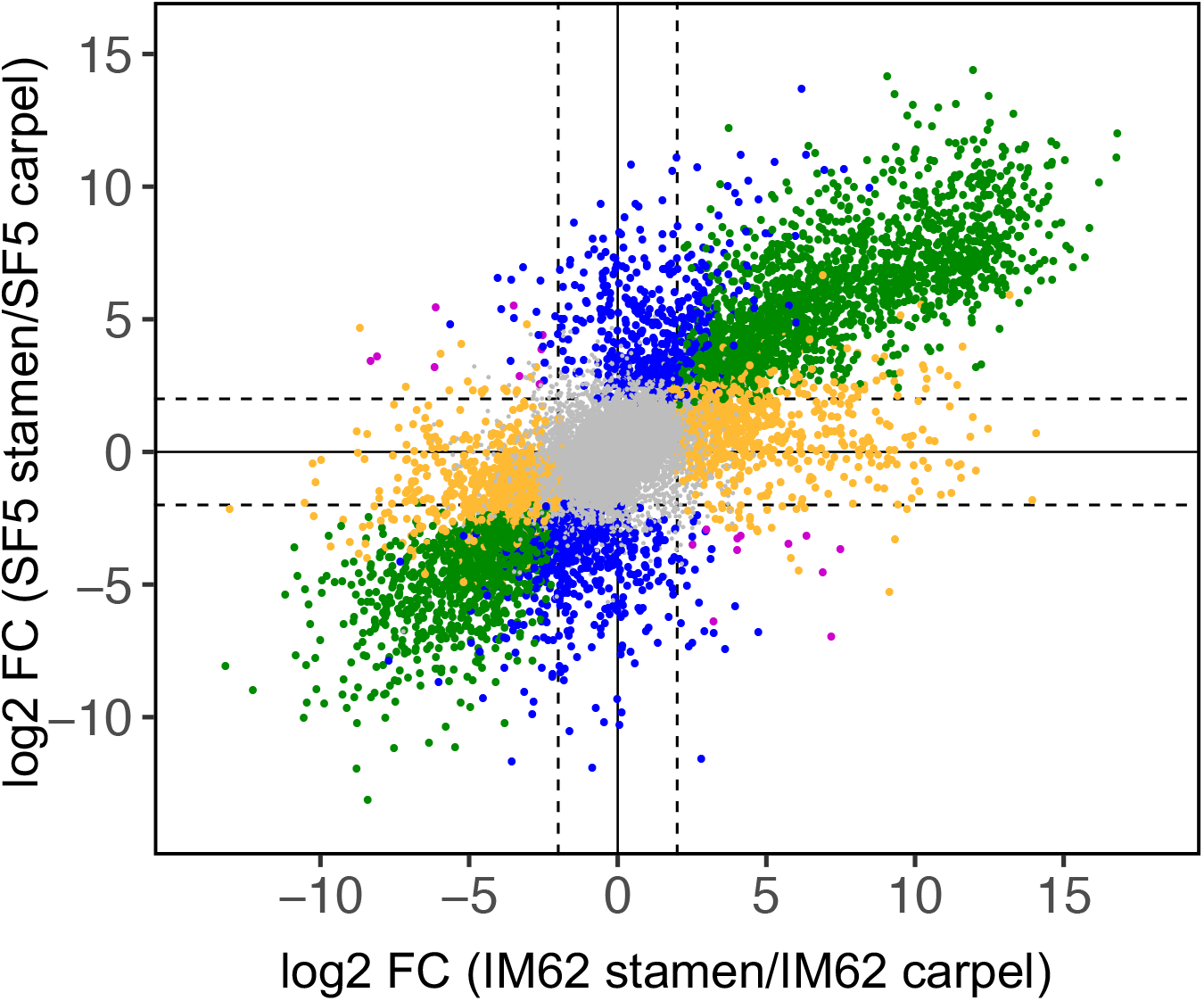
Parental tissue-biased gene expression. Scatterplot shows relative transcript abundance (log2 fold-change (FC)) between stamen and carpel tissues in the SF5 and IM62 parents. Of the 21,147 genes expressed in our dataset, 2038 (9.6%) were stamen-biased (log2 FC>2, FDR≤0.05; top right quadrant green points) and 1355 (6.4%) were carpel-biased (log2 FC<-2, FDR≤0.05; bottom left quadrant green points), in both parents. An additional 598 (2.8%) and 686 (3.2%) genes were stamen- and carpel-biased, respectively, only in SF5 (blue points), while 922 (4.4%) and 1046 (4.9%) genes were stamen- and carpel-biased, respectively, only in IM62 (yellow points). A few genes exhibited opposing tissue-biased expression patterns in SF5 and IM62 (purple points): 9 (0.05%) were stamen-biased in SF5 and carpel-biased in IM62 while 11 (0.05%) were carpel-biased in SF5 and stamen-biased in IM62. The remaining 14,482 (68.5%) genes were evenly expressed between parental tissues (grey points).

A conspicuous exception to the species-tissue clustering pattern just described is the distinct group formed by the STE stamen samples (see red asterisks in Figure 2). We reasoned that this unique pattern of gene expression in STE stamens might be driven by the presence of the chromosome 6 introgression, which contains the hybrid male sterility-causing, IM62 *hms1* allele. To investigate this possibility, we compared gene expression profiles among STE, FER, and SF5 (Figure 4). Indeed, the vast majority of differentially expressed genes were identified in comparisons of stamens with and without the chromosome 6 introgression (STE vs. FER and SF5: 311 upregulated, 1881 downregulated). Moreover, differential expression in STE stamens was pervasive across the genome, affecting roughly equal proportions of genes in introgressed and background regions (Figure S3). Far fewer expression differences were found in comparisons of stamens distinguished only by the chromosome 11 introgression (Figure 4, FER vs. SF5: 9 upregulated, 0 downregulated). Additionally, relatively few differentially expressed genes were found in carpels, whether they carried the chromosome 6 introgression or not (across all comparisons: 27 upregulated, 0 downregulated).

**Figure 4.**
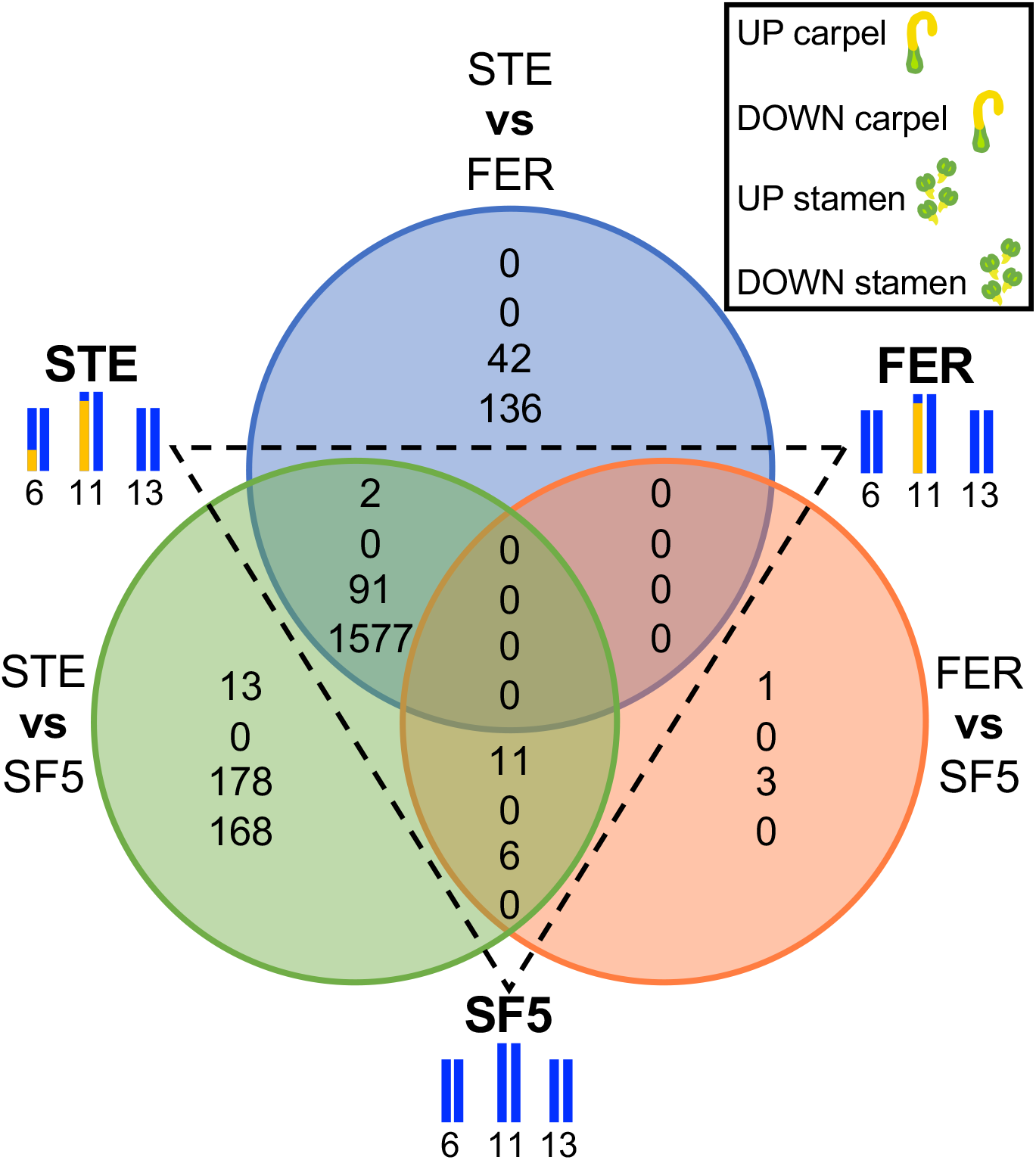
Gene expression differences across sterile (STE), fertile (FER), and SF5 tissues. Venn diagrams show counts of genes with significantly altered transcript abundance (−2<log2 fold-change>2, FDR≤0.05) in carpels and stamens across three pairwise comparisons: (i) RSB_7_ sterile (STE) versus RSB_7_ fertile (FER), (ii) STE versus the recurrent *M. nasutus* SF5 parent, and (iii) FER versus SF5.

### Gene expression in introgression lines is strongly affected by hybrid male sterility

To examine patterns of transcriptional regulation associated with hybrid sterility, we plotted gene expression in FER (Figure 5) and STE (Figure 6) tissues relative to both parents. Because global patterns of relative expression are expected to differ between genes located in introgressions (which carry one IM62 allele and one SF5 allele) and genes located in background regions (which carry two SF5 alleles), we plotted these gene classes separately.

**Figure 5.**
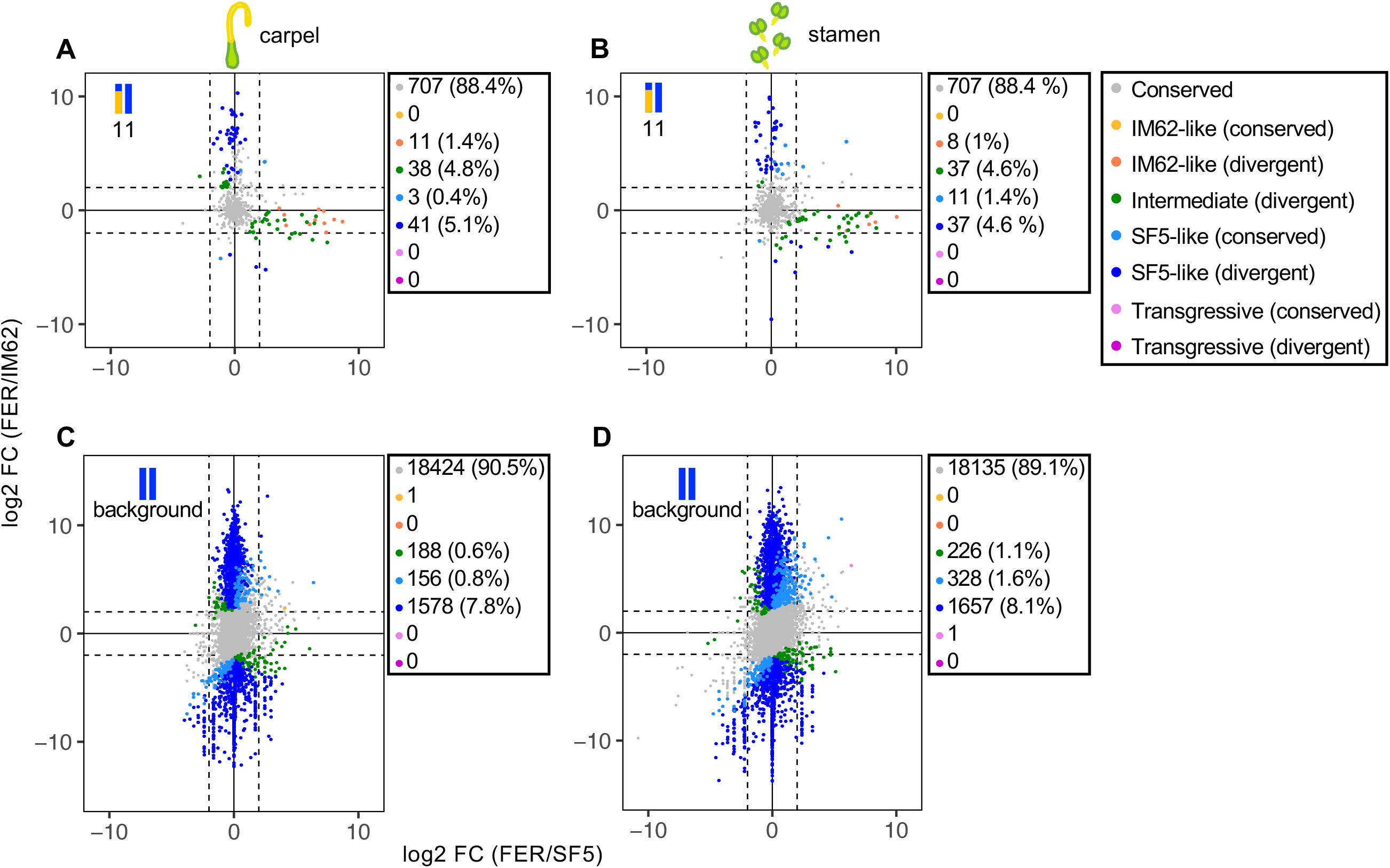
Genome-wide pattern of expression in fertile (FER) hybrids. Plots show relative transcript abundance (log2 fold-change (FC)) between FER and parental (A, C) carpels (B, D) and stamens for (A-B) heterozygous genes in the chromosome 11 introgression and (C-D) homozygous background genes. Genes are colored by expression class (see Table S2 for description).

**Figure 6.**
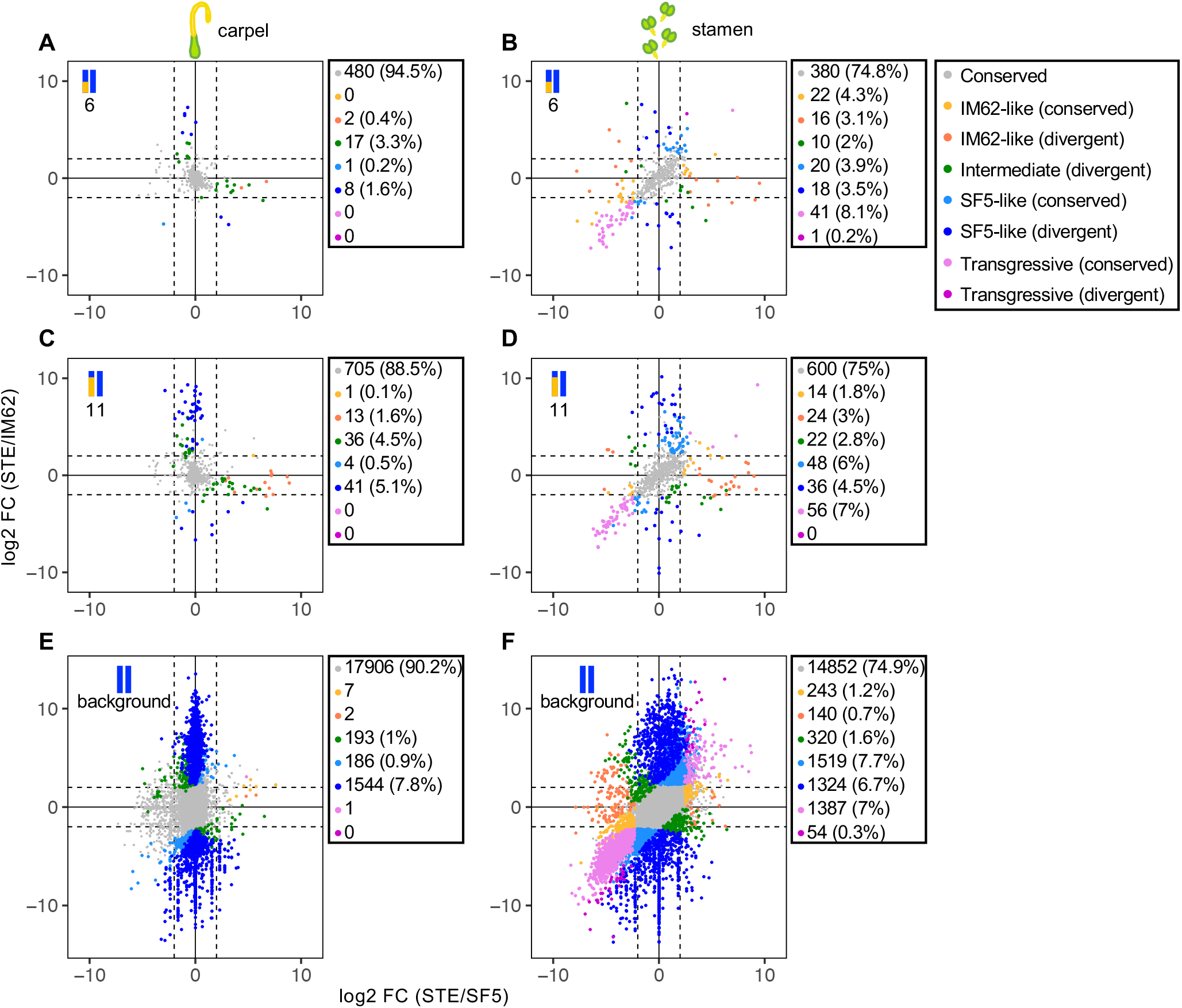
Genome-wide pattern of expression in sterile (STE) hybrids. Plots show relative transcript abundance (log2 fold-change (FC)) between STE and parental (A, C, E) carpels (B, D, F) and stamens for heterozygous genes in the (A-B) chromosome 6 introgression, (C-D) chromosome 11 introgression and (E-F) homozygous background genes. Genes are colored by expression class (see Table S2 for description).

Patterns of relative expression in FER carpels, FER stamens and STE carpels (i.e. tissues unaffected by hybrid sterility) indicate that both *cis*-elements and *trans*-acting factors contribute to regulatory divergence. Among genes that were differentially expressed between the IM62 and SF5 parents (hereafter “IM62-SF5 DEGs”; labeled “divergent” in Figures 5 & 6) and located in either of the two heterozygous introgressions, many showed intermediate levels of expression in FER carpels, FER stamens and STE carpels (green dots in Figures 5A-B and 6A, C), consistent with the additive effects of divergent *cis*-regulatory alleles. Additionally, a substantial number of IM62-SF5 DEGs in the heterozygous introgressions had SF5-like or IM62-like expression levels (dark blue and orange dots in Figures 5A-B and 6A, C), which might result from dominance of one *cis*-acting variant over the other. Consistent with this possibility, expression of heterozygous genes usually matched the more highly expressed parent (note the bias toward positive values in Figures 5A-B and 6A, C). Dominance of a high-expression allele might be achieved via a *cis*-change that increases its own transcription (whereas dominance of a *cis*-acting, low-expression allele would have to *interfere* with expression from the high-expression allele). Effects of *trans*-acting factors on genes in the heterozygous introgressions are also apparent: many more IM62-SF5 DEGs exhibited SF5-like than IM62-like expression levels, suggesting a strong influence of *trans*-factors from the SF5 background (Figures 5A-B and 6A, C). Similarly, although the vast majority of IM62-SF5 DEGs in background regions were SF5-like due to their homozygosity for SF5 alleles (blue dots in Figures 5C-D and 6E), a handful showed IM62-like, intermediate, or transgressive expression (orange, green, and purple dots), consistent with *trans*-acting effects of IM62 alleles from the heterozygous introgressions.

Relative expression in STE stamens differed dramatically from all other samples, with a large number of genes showing transgressive expression (purple dots in Figure 6B, D, F). Additionally, STE stamens showed a marked increase in genes with IM62-like expression, including in background regions where genes are homozygous for SF5 alleles (orange dots in Figure 6B, D, F). Most of these IM62-like background genes were more highly expressed in the SF5 parent (orange dots on the right half of Figure 6F), suggesting they resembled IM62 because of underexpression in STE stamens. Similarly, most transgressive expression was caused by genes that were downregulated relative to both parents (Figure 6F, purple dots: note the bias toward negative values), including many that were not differentially expressed between IM62 and SF5 (light purple dots in Figure 6F).

The fact that transgressive (mostly downregulated) expression was almost entirely restricted to STE stamens suggests an association with the hybrid male sterility phenotype. Consistent with this idea, we observed an overrepresentation of stamen-biased genes among the 2192 genes that were differentially expressed in STE stamens compared to FER and SF5 stamens (see Figure 4): 1372 (63%) were stamen-biased in both parents (74% were stamen-biased in SF5), whereas only 55 (2.5%) were carpel-biased in both parents (5% were carpel-biased in SF5) (Figure S4). The vast majority (1358, 99%) of these 1372 stamen-biased genes were downregulated in STE stamens, whereas most (53, 96%) of the 55 carpel-biased genes were upregulated in this tissue. Moreover, we found overlapping GO term enrichment – mostly relating to pollen tube growth and function – among genes that were downregulated in STE stamens and those that were stamen-biased in the parents (Table S3). Taken together, these results suggest aberrant patterns of gene expression in STE stamens are a consequence of hybrid male sterility.

### Large effects of both *cis-* and *trans-* regulatory divergence between species

Differences in gene expression between species are indicative of divergence in underlying regulatory machinery. By examining allele-specific expression in interspecific hybrids, regulatory divergence can be partitioned among contributing *cis* and *trans* components. Divergence in *cis*-regulatory factors will manifest as biased allele-specific expression in hybrid progeny. In contrast, divergence in *trans-* acting factors affects overall transcript abundance without disrupting allele-specific expression in hybrids. Consequently, a measure of *trans* divergence can be estimated by subtracting the *cis* effects (*i.e.* ratio of allele-specific transcript abundance in hybrids) from the *cis* and *trans* effects (i.e. ratio of transcript abundance between species).

To diagnose the pattern of regulatory divergence in *Mimulus*, we compared expression variation between SF5 and IM62 against allele-specific expression within STE and FER tissues across the heterozygous introgression regions within STE and FER tissues (Figure 7 and S5). Nearly half (31.1-44.5%) of the genes in the introgression regions exhibited both interspecific expression conservation and regulatory conservation (grey dots in Figure 7). Of the genes that were differentially expressed between SF5 and IM62, regulatory divergence was primarily categorized as *trans-* only (21-35, 7.2-11.5% of heterozygous genes; green dots in Figure 7), followed closely by *cis-* only (14-29, 4.8-9.1%; purple dots in Figure 7) and *cis x trans* (10-28, 3.4-7.3% of heterozygous genes; light blue dots in Figure 7). Among genes that were not differentially expressed between SF5 and IM62, many (10-40, 3.7-9.7% of heterozygous genes) showed evidence of compensatory *cis-trans* evolution in FER and STE tissues (orange dots in Figure 7).

**Figure 7.**
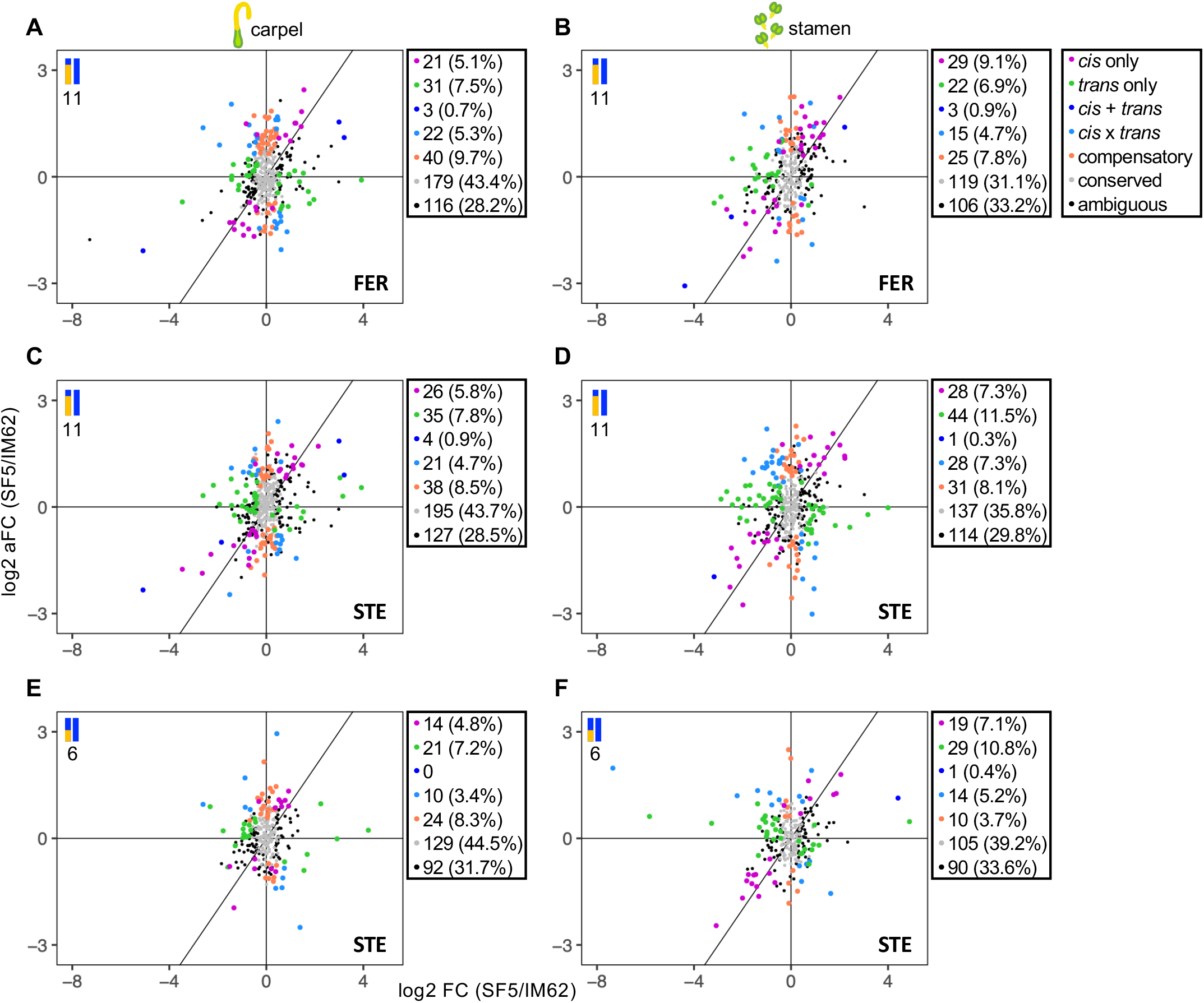
Pattern of *cis-* and *trans-* regulatory differences in FER and STE introgression regions. Plots show relative transcript abundance (log2 fold-change (FC)) between the parents on the x-axis and relative allelic transcript abundance (log2 allelic FC (aFC) within heterozygous introgression regions on the y-axis. Genes are colored by regulatory category.

### Gene expression in *hms1* and *hms2* intervals points to candidate genes

In addition to examining global patterns of gene expression, we wanted to investigate genes in the *hms1*- and *hms2*-mapped intervals to identify candidates for *Mimulus* hybrid sterility. Our expectation is that the causal genes for severe hybrid male sterility and partial hybrid female sterility will be expressed in stamens, and possibly, in carpels. Additionally, if the *hms1-hms2* incompatibility is mediated by expression changes, we might expect the causal genes to show differences in expression between species and/or between fertile and sterile introgression lines.

Of the 11 genes in the *hms1* interval, we were able to evaluate expression for only eight of them. Among these eight genes, six were expressed at moderate to high levels in stamens, with two (Migut.F01606 and Migut.F01607) showing stamen-specificity/bias (Table 1). One of these genes also showed expression differences between species and between introgression lines: Migut.F01606 was highly expressed in SF5 and FER stamens and very lowly expressed (possibly off) in IM62 and STE stamens. One of them (Migut.F01612) shows copy number variation between species: this gene and its highly similar paralog (Migut.F01618) are present in IM62 but absent in SF5. Because of its absence from the genome, expression is precluded in SF5 and FER samples. However, even in IM62 and STE, expression is difficult to gauge because high sequence similarity between Migut.F01612 and Migut.F01618 means that these transcripts did not pass our threshold for unique read mapping. Taken together, these results suggest that Migut.F01606 and Migut.F01612 are the most promising *hms1* candidates.

**Table 1.**
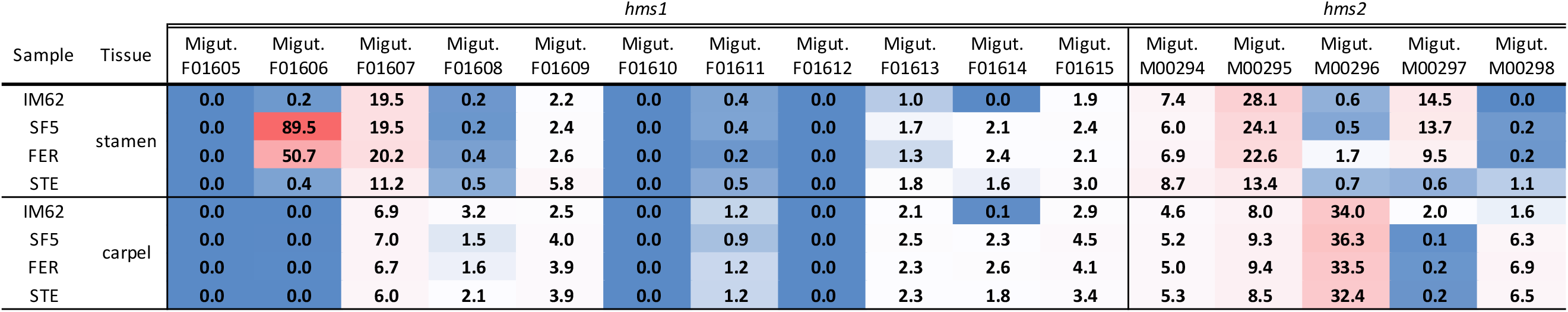
Gene expression at *hms1* and *hms2*. Table shows transcript abundance in fragments per kilobase per million reads sequenced (FPKM) for the 11 and five genes in the mapped regions of *hms1* and *hms2*, respectively.

Of the five genes in the *hms2* interval, three were expressed at moderate to high levels in stamens, with two (Migut.M00295 and Migut.M00297) showing stamen-specificity/bias (Table 1). Two genes (Migut.M00296 and Migut.M00298) were carpel-biased with very little/no expression in stamens, making them unlikely candidates for the *hms2* causal gene. Intriguingly, Migut.M00297 was much more lowly expressed in stamens from STE than from IM62, SF5, or FER. This pattern is precisely what is expected if the IM62 *hms1* allele (in the STE chromosome 6 introgression) directly affects expression of the causal *hms2* gene. From this analysis, then, Migut.M00297 emerges as a strong candidate for *hms2*.

## DISCUSSION

Gene misregulation is a common feature of hybrids between closely related species, but its mechanisms and evolutionary significance are not always clear. Aberrant patterns of gene expression in sterile or inviable hybrids might be due to regulatory incompatibilities – which would implicate divergence in regulatory networks as a driver of reproductive isolation – or to the downstream effects of disrupted tissues. In this study, we took advantage of introgression hybrids between closely related *Mimulus* species to disentangle whether misexpression is a cause or consequence of hybrid sterility. Although we discovered substantial regulatory divergence between *M. guttatus* and *M. nasutus*, regulatory incompatibilities in hybrids were not pervasive. Instead, we found that massive downregulation of global transcript abundance was confined to the affected tissues (*i.e*., stamens) of individuals carrying a sterility-causing introgression on chromosome 6, suggesting that male reproductive development is severely perturbed in sterile hybrids.

Despite the recentness of their split (~200 KYA, Brandvain *et al.*, 2014), *M. guttatus* and *M. nasutus* showed considerable variation in global gene expression, with nearly 10% of genes differentially expressed between species in stamens and/or carpels. This metric likely underestimates the amount of interspecific regulatory divergence in *Mimulus*, both because of our conservatively high threshold for significance (log2 fold-change > 2, equivalent to a four-fold difference in expression) and because of *cis-trans* compensatory evolution, which can lead to changes in underlying regulatory networks while conserving gene expression levels (True & Haag, 2001; Landry *et al.*, 2005). Indeed, eliminating the four-fold threshold cutoff (which we did when categorizing patterns of regulatory divergence), raised the percentage of differentially expressed genes between species to 50%. Extensive expression divergence over short evolutionary timescales has also been observed in animals (Rottscheidt & Harr, 2007; Renaut *et al.*, 2009; Coolon *et al.*, 2014) and other plant systems (Fujimoto *et al.*, 2011; Combes *et al.*, 2015). However, we note that because nucleotide divergence between these *Mimulus* species barely exceeds diversity within *M. guttatus* (*d*_*s*_ = 4.94% and π_*s-M. guttatus*_ = 4.91%, Brandvain *et al.*, 2014), much of the observed expression variation between *M. guttatus* IM62 and *M. nasutus* SF5 might be segregating within species. We also observed more tissue-biased gene expression in *M. guttatus* than in *M. nasutus* (see Figure 3: IM62 has >50% more genes with carpel- and stamen-biased expression). An intriguing possibility is that this difference that might reflect divergence in reproductive traits associated with the species’ distinct mating systems.

Our targeted look at gene expression in two heterozygous introgressions suggests that regulatory differences between *Mimulus* species are often due to divergence in *trans*-factors. This finding runs counter to the expectation that interspecific regulatory variation should be influenced primarily by changes to *cis*-regulatory sequences, which have fewer pleiotropic effects (Wray et al. 2003, Wittkopp et al. 2008). However, it is in agreement with several recent studies showing that *cis*-changes do not always predominate between closely related species (McManus *et al.*, 2010; Meiklejohn *et al.*, 2014; Combes *et al.*, 2015; Guerrero *et al.*, 2016; Metzger *et al.*, 2017). When two lineages have split only recently, some of their regulatory differences might still be polymorphic within species, with purifying selection having had insufficient time to remove deleterious *trans*-regulatory variants. This argument, along with evidence that mutations in *trans*-factors arise more frequently (Landry *et al.*, 2007), might explain the higher contribution of *trans*-acting factors to regulatory variation observed within species (Wittkopp *et al.*, 2008; Emerson *et al.*, 2010), as well as between closely related species. In the case of the two *Mimulus* species studied here, an additional factor may influence the predominance of *trans*-regulatory divergence: deleterious mutations might be particularly likely to accumulate in *M. nasutus* because of its shift to self-fertilization. Indeed, this species shows genomic signatures that indicate a reduction in the efficacy of purifying selection (Brandvain *et al.*, 2014), an expected outcome of the lower effective population size and recombination rate that accompanies the evolution of selfing (Nordborg, 2000; Charlesworth & Wright, 2001).

In addition to *cis*- and *trans*-only regulatory divergence between these *Mimulus* species, our analyses of the two heterozygous introgressions uncovered evidence of *cis* x *trans* compensatory evolution. Interestingly, however, we found little indication that this process acts as a general driver of regulatory incompatibilities. Genes defined as compensatory were no more likely to be misexpressed (i.e., expressed outside the range of IM62 or SF5) than genes with conserved regulation (Figure S5). This result is somewhat surprising because compensatory changes within species are expected to cause mismatches in hybrids between *cis*- and *trans*-regulators, leading to aberrant gene expression and, potentially, hybrid dysfunction (Landry *et al.*, 2005). In support of this idea, several studies have shown that genes with *cis*- and *trans*-variants that act in opposing directions (i.e., *cis* x *trans*) are enriched among genes that are misexpressed in hybrids (Tirosh *et al.*, 2009; McManus *et al.*, 2010; Schaefke *et al.*, 2013; Mack *et al.*, 2016). On the other hand, it has been argued that *cis* x *trans* effects might often be inflated (Fraser, 2019), and, in some cases, the association is missing altogether (Bell *et al.*, 2013; Coolon *et al.*, 2014).

What is clear from our study is that widespread misexpression in *Mimulus* introgression hybrids is caused not by regulatory divergence, but by the hybrid sterility phenotype itself. Introgressing a 7 Mb genomic segment with the *hms1* incompatibility allele from *M. guttatus* into *M. nasutus* has profound effects on male fertility, with RSB_7_ STE individuals producing only few pollen grains, nearly all of which are inviable. Coincident with this male sterility, RSB_7_ STE stamens showed dramatic expression differences when compared to parental lines, with 7.2% of all genes misexpressed (*N* = 21,147). In stark contrast, introgressing a genomic segment from chromosome 11, which is much larger in size (23 Mb) but does not carry any known hybrid incompatibility alleles, resulted in only a single misexpressed transcript in stamens (i.e., in RSB_7_ FER individuals). The fact that neither introgression showed strong effects on expression in carpels suggests that partial hybrid female sterility either has few effects on gene regulation or manifests later in development.

Our study highlights the challenge of distinguishing the potential regulatory causes of hybrid incompatibilities from downstream effects. In subspecies of house mice, seven *trans*-eQTL colocalize with QTL for hybrid sterility (Turner *et al.*, 2014), providing strong evidence for a causal link between divergent regulatory alleles and the evolution of hybrid incompatibilities. In most other cases, however, this link has been difficult to establish because when hybrid misexpression is discovered, it is often confounded with the phenotypic effects of hybrid dysfunction, such as defective tissues and/or disrupted development (Ortíz-Barrientos *et al.*, 2007; Wei *et al.*, 2014). For example, when normally inviable F1 hybrid males between *Drosophila melanogaster* and *D. simulans* are rescued by a mutation in the hybrid incompatibility gene *Hmr*, gene expression in larvae becomes much more similar to parents (Wei *et al.*, 2014). Like with the *hms1-hms2* incompatibility in *Mimulus*, this result suggests a large effect of the *Hmr* gene on genome-wide hybrid misexpression in *Drosophila*. A similar result was also seen in a previous study of *M. nasutus-M. guttatus* F2 hybrids: the number of differentially expressed genes between parents and lethal F2 seedlings, which lack chlorophyll due to a two-locus hybrid incompatibility, is much higher than between parents and viable F2 seedlings (Zuellig & Sweigart, 2018). Taken together, these studies suggest that caution is needed when assigning a cause to hybrid gene misexpression. At the same time, it is important to note that our results do not rule out regulatory divergence as a cause of *Mimulus* hybrid incompatibilities in particular cases.

In addition to the main findings just discussed, our study has revealed a dramatic suppression of recombination in the RSB introgression population. Despite eight rounds of backcrossing to *M. nasutus*, the heterozygous introgressions on chromosomes 6 and 11 remain quite large (7 Mb and 23 Mb, respectively). With uniform recombination rates and Mendelian transmission, RSB_7_ individuals are expected to be heterozygous along ~0.2% of their genome, which equates to a maximum introgression size of ~0.625 Mb (*M. guttatus* genome ~312 Mb). Suppressed recombination rates on chromosome 6 were observed previously, in an earlier generation of the RSB population (Sweigart *et al.*, 2006). At the time, we speculated that low recombination might be a direct cause of the *hms1-hms2* incompatibility – perhaps due to a meiotic defect. However, follow-up work performing testcrosses with F2 hybrids that carried either incompatible or parental genotypes at *hms1* and *hms2* showed no effect of the *hms1-hms2* incompatibility on recombination rates (data not shown). Additionally, we have observed a similar reduction in recombination in heterospecific introgressions when attempting to generate nearly isogenic lines using other *Mimulus* accessions that lack the *hms1-hms2* incompatibility. An alternative explanation for the suppressed recombination on chromosomes 6 and 11 is that local sequence diversity affects recombination in *Mimulus*. In our crossing scheme, nucleotide diversity between chromosome homologs was much higher in heterospecific introgressions than in adjacent isogenic regions. Thus, if sequence diversity affects the likelihood of DNA double-strand breaks and/or crossover events, as it does in mice (Li *et al.*, 2018), we would expect much lower recombination in the heterozygous introgressions. Given the extraordinarily high nucleotide diversity within *M. guttatus* (Brandvain *et al.*, 2014; Puzey *et al.*, 2017), if this explanation is correct, we might expect extensive natural variation in recombination rates even within species.

Finally, we note that our study has shortened the list of likely candidate genes for causing the *hms1-hms2* incompatibility. Previously, we mapped *hms1* to an interval containing 11 annotated genes with three strong functional candidates: *Migut.F01605*, *Migut.F01606*, and *Migut.F01612* (Sweigart & Flagel, 2015). The first two are tandem duplicates of *SKP1*-like genes, which form part of the SKP1–Cullin–F-box protein (SCF) E3 ubiquitin ligase complex that regulates many developmental processes including the cell cycle (Hellmann and Estelle 2002). Although we did not detect *Migut.F01605* expression in any sample (potentially calling into question its functionality), *Migut.F01606* remains a strong candidate. Expression of this gene was stamen-specific and much higher in *M. nasutus* SF5 and RSB_7_ FER than in *M. guttatus* IM62 or RSB_7_ STE. If *Migut.F01606* is causal for *hms1*, the fact that the normally expressed SF5 allele is off in heterozygous RSB_7_ STE individuals suggests that the IM62 allele interferes with its expression. An alternative possibility is that reduced expression of *Migut.F01606* in RSB_7_ STE is only one of the many downstream effects of the hybrid male sterility phenotype. *Migut.F01612*, an F-box gene, also remains a strong candidate for *hms1*. Although the RNAseq results provided little additional insight into the function of this gene (we did not detect expression in any sample), we have observed its expression in IM62 via RT-PCR (data not shown) and its absence from the SF5 genome is notable.

At *hms2*, expression patterns of *Migut.M00297* strengthen it as a candidate. This gene encodes the second-largest subunit (*RPB2*) of RNA Polymerase II – the multi-subunit enzyme responsible for mRNA transcription (Woychik & Hampsey, 2002; Hahn, 2004). In most flowering plant species, *RPB2* is a single copy gene. However, in the asterid clade, two distinct paralogs (*RPB2*-*i* and *RPB2*-*d*) are present, having been retained following an ancient duplication event (Oxelman *et al.*, 2004; Luo *et al.*, 2007). In all asterid species that have been investigated, the expression pattern of *RPB2-i* suggests that it is restricted to male reproductive structures (e.g. stamen and pollen) (Oxelman *et al.*, 2004; Luo *et al.*, 2007). In our experiment, expression of *Migut.M00297*, which encodes the *RPB2-i* paralog, was highly stamen-biased in both parents and in RSB_7_ FER, but off in RSB_7_ STE individuals. Although this is the pattern expected if the IM62 *hms1* allele (in the STE chromosome 6 introgression) directly affects the expression of the causal *hms2* gene, it might also arise as a byproduct of *hms1-hms2* sterility. Of course, an important consideration is that, for both *hms1* and *hms2*, the difference between compatible and incompatible alleles might have nothing to do with transcription. For each of these loci, then, additional approaches such as transformation experiments will be needed to identify the causal genes.

## METHODS

### Plant lines and growth conditions

This study focuses on *Mimulus guttatus* and *M. nasutus*, two closely related species that diverged roughly 200,000 years ago (Brandvain *et al.*, 2014). Previous work identified two nuclear incompatibility loci – *hms1* and *hms2* – that cause nearly complete male sterility and partial female sterility in a fraction of F_2_ hybrids between an inbred line of *M. guttatus* from Iron Mountain, Oregon (IM62), and a naturally inbred *M. nasutus* line from Sherar’s Falls, Oregon (SF5). We generated an introgression population carrying incompatible (IM62) and compatible (SF5) *hms1* alleles in a common genetic background through multiple rounds of selection for pollen sterility and backcrossing to the recurrent SF5 parent (Sweigart *et al*. 2006). Briefly, *M. nasutus* SF5 and *M. guttatus* IM62 were intercrossed (with SF5 as the maternal parent) to create an F_1_ hybrid that was backcrossed to SF5 (with the F1 hybrid as the maternal parent). This crossing scheme resulted in a first-generation backcross (BC_1_) population that segregates four *hms1-hms2* genotypes (Figure 1). Next, a pollen sterile individual selected from the BC_1_ population was backcrossed SF5, yielding the first generation of an introgression population dubbed *r*ecurrent *s*election with *b*ackcrossing (i.e. RSB_1_) (Sweigart *et al.*, 2006). This selective backcrossing scheme was repeated for six more generations to produce an RSB_7_ population that segregates approximately 1:1 for two genotypes: pollen sterile (STE) individuals carrying a heterozygous introgression of the incompatible IM62 allele at *hms1* and pollen fertile (FER) individuals homozygous for the compatible SF5 allele at *hms1*, both in a genetic background that is fixed for the incompatible SF5 allele at *hms2*. To identify FER and STE RSB_7_ plants prior to flowering, individuals were genotyped at markers flanking *hms1* and *hms2*. To verify the fertility phenotypes, the first flower on each RSB_7_ plant was allowed to self-pollinate. Within three to five days post-anthesis, fertilized fruits (on FER plants) begin to mature and plump, while the unfertilized fruits (on STE plants), remain immature and small, making it easy to differentiate the two phenotypic classes.

All plants were grown in a growth chamber at the University of Georgia. Seeds were sown into 2.5-inch pots containing Fafard 3B potting mix (Sun Gro Horticulture, Agawam, MA), stratified for 7 days at 4°C, then transferred to a growth chamber set to 22C day/16C night, 16-hour day length. Plants were bottom-watered daily and fertilized with bloom booster as needed.

### Sample collection and transcriptome sequencing

For this study, 24 whole transcriptome libraries were generated, consisting of three bioreps each of two tissue types (i.e. stamens and carpels) from four genotypes that vary at *hms1* and *hms2* (i.e. *M. guttatus* IM62, *M. nasutus* SF5, RSB_7_ fertile [FER], and RSB_7_ sterile [STE]) (Figure 1, and Table S1). To collect enough tissue for each biorep, we carefully dissected 8-24 pre-anthesis floral buds and transferred the carpels and stamens to 1.5-mL microcentrifuge tubes that were partially submerged in liquid nitrogen (Table S1). For each sample, we extracted RNA using a QuickRNA Miniprep Kit (Zymo Research, Irvine, CA, USA) then assayed and measured RNA concentration using a Qubit RNA BR (Broad-Range) Assay Kit and a Qubit 2.0 Fluorometer (Thermo Fisher Scientific Inc., Waltham, MA, USA). Samples were shipped overnight on dry ice to the Duke Center for Genomic and Computational Biology (Durham, NC, USA), where the Sequencing and Genomic Technologies core checked RNA quality using an Bioanalyzer 2100 (Agilent Technologies, Santa Clara, CA, USA), constructed sequencing libraries using a KAPA Stranded mRNA-Seq Kit (F. Hoffmann-L Roche, Basel, Switzerland), and sequenced all 24 libraries on a single lane of HiSeq 4000 (Illumina, Inc. San Diego, CA, USA), producing 50-base pair (bp) single-end reads (Table S1).

### *M. nasutus* pseudoreference genome construction and transcriptome alignment

An important consideration for genomic and transcriptomic analyses is potential mapping bias introduced when aligning reads from one species against a reference genome or transcriptome from another species (Degner *et al.*, 2009). The four genotypes in our dataset IM62, SF5, FER and STE, represent two pure species, *M. guttatus* and *M. nasutus*, as well as fertile (FER) and sterile (STE) siblings from a *M. nasutus*-*M. guttatus* backcrossed line. Previous work showed that interspecific nucleotide divergence between *M. guttatus* and *M. nasutus* is substantial (*d*_*S*_ = 4.94%, see Brandvain et al. 2014). Therefore, aligning SF5, FER and STE – all of which are expected to carry SF5 alleles across >90% of their genomes – against the *M. guttatus* v2.0 reference genome (Hellsten *et al.*, 2013) is likely to cause mapping bias due to mismatch errors. Reads originating from *M. nasutus* may align incorrectly, non-uniquely, or not at all against the *M. guttatus* v2.0 reference genome. Further magnifying potential mapping bias issues, the *M. guttatus* v2.0 reference genome assembly is based on the IM62 accession and as such, IM62-derived reads are expected to align with near perfection. To ameliorate this issue, we first constructed a *M. nasutus* pseudoreference genome using publicly available SF5 whole genome sequence data, then aligned our SF5, FER and STE RNAseq reads against this.

Using the fastq-dump command from the NCBI toolkit, we retrieved the SF5 gDNA fastq files from the NCBI SRA database (SRR400478). To prepare these 75-bp paired-end sequences for alignment, we trimmed adapters and low-quality bases, then filtered out processed reads shorter than 50 bp using Trimmomatic (Bolger *et al.*, 2014). We mapped the resulting 50-75-bp paired-end reads to the *M. guttatus* v2.0 reference genome using BWA-MEM (Li & Durbin, 2009; Li, 2013). To filter the initial SF5 alignment, we eliminated optical and PCR duplicates using the MarkDuplicates tool from Picard (http://broadinstitute.github.io/picard) and removed reads with an alignment quality below Q30 using the view command from SAMtools (Li *et al.*, 2009). Next, we generated a set of high-quality SF5 single nucleotide polymorphisms (SNPs) to use in pseudoreference genome construction. First, we used GATK’s HaplotypeCaller in GVCF mode followed by GATK’s GenotypeGVCFs to identify phased SNP and insertion/deletion (indel) variants in the SF5 alignment (McKenna *et al.*, 2010; Poplin *et al.*, 2017). Next, we extracted biallelic SNPs using GATK’s SelectVariants tool and filtered out sites with a mapping quality (MQ) below 40 or quality by depth (QD) below two using GATK’s VariantFiltration. We used this filtered biallelic SNP VCF file to generate a *M. nasutus* pseudoreference using GATK’s FastaAlternateReferenceMaker tool.

To increase mapping fidelity, we aligned our RNAseq reads against the appropriate reference: samples from the IM62 parent were aligned to the *M. guttatus* v2.0 reference genome and all other samples were aligned to the *M. nasutus* pseudoreference. Before mapping, we trimmed adapter sequences and low-quality bases, then filtered out processed reads shorter than 36 bp using Trimmomatic (Bolger *et al.*, 2014) (Table S1). The resulting 36-50-bp single-end RNAseq reads were aligned to the *M. guttatus* v2.0 reference genome with STAR in the multi-sample 2-pass mapping mode (Dobin *et al.*, 2013; Dobin & Gingeras, 2015). Transcriptome alignments were filtered similarly to genome alignments, with one additional step. We removed optical and PCR duplicates with the MarkDuplicates command from Picard (http://broadinstitute.github.io/picard), then used the SplitNCigarReads command from GATK to parse intron-spanning reads into exon segments and trim bases extending into intronic regions (McKenna *et al.*, 2010). Finally, we removed reads with an alignment quality below Q30 using the view command from SAMtools (Li *et al.*, 2009).

### Variant calling and identification of introgression boundaries in RSB samples

We used the GATK HaplotypeCaller tool in GVCF mode to call single nucleotide polymorphisms (SNPs) and insertions/deletions (indels) in the processed RNAseq alignment files. Variant calling from transcriptome data is dependent on transcript abundance, which can vary across genotypes and tissues. To increase our power to call variants in our 24 RNAseq samples, we simultaneously genotyped them alongside SF5 and IM62 DNAseq samples (SRR400478 and SRR052268) with the GATK GenotypeGVCF tool. Following joint genotyping, we performed a series of filtering steps using GATK and bcftools. First, sites with a mapping quality (MQ) score below 30 or quality by depth (QD) below two were filtered from the multi-sample variant call file (VCF). Then, for each sample, VCF files containing only biallelic SNPs were extracted from the multi-sample VCF and filtered individually. Next, sites with a read depth below five were excluded from individual samples. Finally, sites were excluded from all samples if they were (i) heterozygous in the IM62 or SF5 parental samples, (ii) homozygous reference in the SF5 samples (i.e. SF5 = IM62 reference), or (iii) homozygous non-reference in the IM62 samples (i.e. IM62 ≠ IM62 reference), as these were confounding or uninformative. After these filtering steps, we retained variant sites called in at least 50% (12/24) of the RNAseq samples, resulting in 198,072 high-confidence SNPs. We extracted chromosomal location and genotype information at each SNP for all 24 RNAseq samples with bcftools query, which we used to identify introgression boundaries in the RSB samples.

### Differential gene expression analysis

To estimate transcript abundance, reads were counted in the final processed transcriptome alignment files using HTSeq (Anders *et al.*, 2015). These raw read counts were then utilized to perform differential gene expression in edgeR (Robinson *et al.*, 2010). To restrict comparisons to genes expressed in at least one genotype-by-tissue group, we included only genes with at least one read count-per-million (CPM) in three or more of the 24 libraries. This filtering step removed 6,361 genes, resulting in a set of 21,779 expressed genes. The calcNormFactors function was used normalize libraries for RNA composition with the default trimmed mean of M-values (TMM) method. To obtain a global view of gene expression across the 24 samples in our dataset, we used the plotMDS function in edgeR to generate a multidimensional scaling (MDS) plot, which is a type of unsupervised clustering plot. The distance between two points in an MDS plot represents the leading log-fold-change (i.e. largest absolute log-fold-change) between that sample pair. To test for differences in gene expression across the genotype-by-tissue groups, we conducted generalized linear model (GLM) analyses using a quasi-likelihood (QL) approach in edgeR. This method is flexible and permits any combination of sample comparisons to be made. First, we generated an experimental design matrix describing the eight genotype-by-tissue groups using the model.matrix function, then fitted it to a quasi-likelihood GLM framework using the glmQLFit function in edgeR. To identify genes for which the log2 fold-change (log2 FC) was significantly greater than two for a given comparison, we used the glmTreat function in edgeR, which performs threshold hypothesis testing on the GLM specified by the glmQLFit function. This is a rigorous statistical test that detects expression differences greater than the specified threshold value by evaluating both variability and magnitude of change in expression, then applies false discovery rate (FDR) *p*-value corrections. We categorized genes as significantly differentially expressed between two genotype-by-tissue groups if the log2 FC in transcript abundance was greater than two and the FDR-corrected *p*-value was less than or equal to 0.05.

### Gene expression category assignment

We categorized gene expression in the STE and FER RSB_7_ siblings based on interspecific expression differences between IM62 and SF5, as well as differences between the RSB individual and its two parents (Table S2). This resulted in the following eight categories: (i) Conserved: the parents and RSB all have similar expression; (ii) SF5-like-conserved: RSB expression is similar to SF5 and significantly different than IM62. Expression does not differ between parents; (iii) SF5-like-divergent: RSB expression is similar to SF5 and significantly different from IM62. Expression differs significantly between parents; (iv) IM62-like-conserved: RSB expression is similar to IM62 and significantly different from SF5. Expression does not differ between parents; (v) IM62-like-divergent: RSB expression is similar to IM62 and significantly different from SF5. Parents differ significantly; (vi) Intermediate: RSB expression falls within the parental range. Expression differs significantly between parents; (vii) Transgressive-conserved: RSB expression is higher or lower than both parents. Expression does not differ between parents; (viii) Transgressive-divergent: RSB expression is higher or lower than both parents. Expression differs significantly between parents.

### Allele-specific expression analysis

To measure allele-specific expression (ASE) within the heterozygous regions of FER and STE RSB_7_ siblings, we used the phASER tool suite (Castel *et al.*, 2016). Only reads that overlap polymorphic sites are useful for ASE estimation. We quantified allele-specific counts across individual variant sites within the heterozygous regions of FER and STE samples using the phASER tool. We limited our ASE quantification to the 198,072 high-confidence SNP sites we previously identified (described above). As a filtering step, we removed allele-specific counts at sites with a read depth below five in individual samples. Next, we produced gene-level allele-specific counts at each heterozygous gene by summing counts across SNP sites located in the same gene with the phaser Gene AE tool. We utilized these gene-level allele-specific counts to perform ASE analyses in edgeR (Robinson *et al.*, 2010). We restricted analyses to genes located in the chromosome 6 and chromosome 11 introgressions with one or more allele-specific CPM in at least at least one genotype-by-tissue group. These filtering steps removed 391 of the 1308 genes expressed in the introgression regions, leaving 917 for ASE analysis. To test for differences in ASE across FER and STE tissues, we fitted our allele-specific count data to a GLM then performed likelihood ratio tests using the glmFit and glmLRT functions in edgeR. We categorized genes as having significant allelic imbalance within a genotype-by-tissue group if the log2 transformed ratio of allele-specific transcript abundance was greater than zero and the FDR-corrected *p*-value was less than or equal to 0.1.

### *Cis-*and *trans-* regulatory divergence category assignment

To estimate total (*cis* and *trans*) interspecific regulatory divergence in *Mimulus*, the log2 transformed ratio of transcript abundance between SF5 and IM62 (pFC) was calculated across stamens and carpels. To estimate *cis-* regulatory divergence, the log2 transformed ratio of allele-specific transcript abundance (aFC) was computed for each heterozygous gene in the chromosome 6 and chromosome 11 introgression regions across FER and STE carpels and stamens. To estimate *trans* effects (*trans*) across FER and STE carpels and stamens, aFC was subtracted from the pFC in the corresponding tissue. To test for regulatory divergence, we analyzed pFC and aFC across FER and STE carpels and stamens for heterozygous genes in the introgression regions. For the purpose of regulatory divergence categorization, pFC tests were performed using glmQLFTest() function in edgeR, eliminating the four-fold expression threshold used in previous analyses. Significant difference in parental gene expression (*i.e.* significant pFC) was considered evidence of total (*cis* and *trans*) regulatory divergence. Similarly, significant imbalance in allelic ratio (*i.e.* significant aFC) in the introgression hybrids was considered evidence of *cis-* regulatory divergence. Genes with significant pFC or significant aFC were analyzed for significant *trans* effects by comparing pFC to aFC using *Student’s t-test*. Significant differences (*p*-value <= 0.1) between these two ratios was considered evidence for *trans-* divergence. Using these results, we partitioned regulatory divergence across FER and STE carpels and stamens into the following seven categories: (i) *Cis* only: Significant pFC and aFC. Non-significant *trans.* The magnitude of aFC is greater than the magnitude of *trans*. pFC and aFC have the same sign (*i.e.* the species with higher expression also had the higher expressing *cis-* allele); (ii) *Trans* only: Significant pFC and *trans*. Non-significant aFC. The magnitude of *trans* is greater than the magnitude of aFC. pFC and *trans* have the same sign (i.e. the species with higher expression also had the higher expressing *trans*-allele); (iii) *Cis + trans:* Significant pFC, aFC, and *trans*. aFC and *trans* have the same sign (*i.e.* the species with higher expressing *cis-* allele also had the higher expressing *trans*-allele); (iv) *Cis* x *trans*: Significant pFC, aFC, and *trans*. aFC and *trans* have the opposite sign (*i.e.* the species with higher expressing *cis-* allele had the lower expressing *trans*-allele and vice versa); (v) Compensatory: Non-significant pFC. Significant aFC, and *trans*. aFC and *trans* have the opposite sign (*i.e.* the species with higher expressing *cis-* allele had the lower expressing *trans*-allele and vice versa); (vi) Conserved: Non-significant pFC and aFC; (vii) Ambiguous: Any other combination of pFC, aFC, and *trans* (these have no clear interpretation).

### Gene Ontology enrichment analysis

We performed Gene Ontology (GO) term analysis using the PlantRegMap online server (http://plantregmap.cbi.pku.edu.cn/index.php). To identify overrepresented GO terms within sets of differentially expressed genes, a significance threshold of a p-value ≤ 0.01 was chosen.

## ACKNOWLEDGMENTS

The Sequencing and Genomics Technology core at the Duke Center for Genomic and Computational Biology prepared RNA sequencing libraries and performed whole transcriptome sequencing. Computational analyses associated with transcriptome alignment, variant detection, and total and allele-specific read quantification were conducted on the Sapelo/Sapelo2 high-performance computing cluster maintained by the Georgia Advance Computing Resource Center (GACRC) at UGA. We thank Casey Bergman, Kelly Dyer, and Dave Hall for valuable discussions. We also thank Samuel Mantel, Colin Meiklejohn, Gabrielle Sandstedt, and Matthew Zuellig for thoughtful comments on an earlier draft, which greatly improved the manuscript. This work was supported by National Science Foundation grant DEB-1350935 to ALS. Additionally, REK was supported by National Science Foundation grant IOS-PGRP-1546617.

