## Supplemental Materials for "Genome-wide misexpression associated with hybrid sterility in *Mimulus* (monkeyflower)"

**Table S1. Transcriptome library sample, sequencing and alignment details.** Table show information for each of the 24 samples in this study. *M. guttatus* = *Mimulus guttatus;* IM62 = parent inbred line from Iron Mountain, Oregon. *M. nasutus* = *Mimulus nasutus* ; SF5 = parent inbred line from Sherar's Falls, Oregon; RSB7 = SF5-IM62 *r* ecurrent *s* election with *b* ackcrossing introgression line (7th inbred generation); FER = fertile RSB7 genotypes; STE = sterile RSB7 genotypes.

| Sample | Genotype | Tissue | Group | Biorep | # buds pooled | RNA conc (ng/uL) | Raw read count | Trimmed read count | % reads trimmed | Reference genome | % uniquely mapped | % multi mapped | % mismatch |
|---|---|---|---|---|---|---|---|---|---|---|---|---|---|
| RK01 | *M. guttatus* IM62 | stamen | IM62st | 1 | 8 | 468 | 13,382,768 | 13,338,218 | 0.33 | *M. guttatus* v2.0 | 88.3 | 5.3 | 0.2 |
| RK02 | *M. guttatus* IM62 | stamen | IM62st | 2 | 8 | 326 | 12,913,305 | 12,869,163 | 0.34 | *M. guttatus* v2.0 | 85.9 | 5.0 | 0.2 |
| RK03 | *M. guttatus* IM62 | stamen | IM62st | 3 | 8 | 457 | 13,105,458 | 13,058,526 | 0.36 | *M. guttatus* v2.0 | 85.5 | 5.2 | 0.2 |
| RK04 | *M. guttatus* IM62 | carpel | IM62cp | 1 | 8 | 93 | 13,865,825 | 13,809,569 | 0.41 | *M. guttatus* v2.0 | 64.5 | 3.7 | 0.2 |
| RK05 | *M. guttatus* IM62 | carpel | IM62cp | 2 | 8 | 342 | 12,758,027 | 12,714,667 | 0.34 | *M. guttatus* v2.0 | 78.6 | 4.0 | 0.2 |
| RK06 | *M. guttatus* IM62 | carpel | IM62cp | 3 | 8 | 382 | 14,922,318 | 14,872,515 | 0.33 | *M. guttatus* v2.0 | 68.9 | 4.0 | 0.2 |
| RK07 | *M. nasutus* SF5 | stamen | SF5st | 1 | 16 | 398 | 16,789,369 | 16,729,667 | 0.36 | *M. nasutus* pseudoreference | 85.1 | 5.1 | 0.3 |
| RK08 | *M. nasutus* SF5 | stamen | SF5st | 2 | 16 | 107 | 10,920,821 | 10,883,494 | 0.34 | *M. nasutus* pseudoreference | 83.8 | 5.0 | 0.3 |
| RK09 | *M. nasutus* SF5 | stamen | SF5st | 3 | 16 | 361 | 14,287,104 | 14,235,966 | 0.36 | *M. nasutus* pseudoreference | 84.3 | 5.0 | 0.3 |
| RK10 | *M. nasutus* SF5 | carpel | SF5cp | 1 | 16 | 122 | 13,021,522 | 12,977,343 | 0.34 | *M. nasutus* pseudoreference | 72.7 | 3.9 | 0.2 |
| RK11 | *M. nasutus* SF5 | carpel | SF5cp | 2 | 16 | 399 | 13,672,385 | 13,626,516 | 0.34 | *M. nasutus* pseudoreference | 77.3 | 4.1 | 0.2 |
| RK12 | *M. nasutus* SF5 | carpel | SF5cp | 3 | 16 | 600 | 16,728,757 | 16,674,470 | 0.32 | *M. nasutus* pseudoreference | 68.4 | 3.7 | 0.2 |
| RK13 | $RSB_7$ FER | stamen | FERst | 1 | 16 | 151 | 16,780,282 | 16,723,300 | 0.34 | *M. nasutus* pseudoreference | 72.9 | 4.7 | 0.3 |
| RK14 | $RSB_7$ FER | stamen | FERst | 2 | 16 | 80 | 16,800,373 | 16,743,146 | 0.34 | *M. nasutus* pseudoreference | 73.5 | 4.5 | 0.3 |
| RK15 | $RSB_7$ FER | stamen | FERst | 3 | 16 | 130 | 14,540,795 | 14,489,407 | 0.35 | *M. nasutus* pseudoreference | 73.2 | 4.6 | 0.3 |
| RK16 | $RSB_7$ FER | carpel | FERcp | 1 | 16 | 288 | 12,592,991 | 12,550,331 | 0.34 | *M. nasutus* pseudoreference | 70.6 | 4.0 | 0.3 |
| RK17 | $RSB_7$ FER | carpel | FERcp | 2 | 16 | 307 | 11,813,346 | 11,774,678 | 0.33 | *M. nasutus* pseudoreference | 77.5 | 4.0 | 0.3 |
| RK18 | $RSB_7$ FER | carpel | FERcp | 3 | 16 | 292 | 12,445,173 | 12,402,433 | 0.34 | *M. nasutus* pseudoreference | 81.6 | 4.1 | 0.3 |
| RK19 | $RSB_7$ STE | stamen | STEst | 1 | 24 | 49 | 13,998,392 | 13,952,091 | 0.33 | *M. nasutus* pseudoreference | 77.3 | 3.5 | 0.3 |
| RK20 | $RSB_7$ STE | stamen | STEst | 2 | 24 | 47 | 13,216,235 | 13,170,400 | 0.35 | *M. nasutus* pseudoreference | 79.9 | 3.7 | 0.3 |
| RK21 | $RSB_7$ STE | stamen | STEst | 3 | 24 | 43 | 14,960,626 | 14,910,714 | 0.33 | *M. nasutus* pseudoreference | 79.2 | 3.6 | 0.3 |
| RK22 | $RSB_7$ STE | carpel | STEcp | 1 | 16 | 148 | 13,427,383 | 13,382,886 | 0.33 | *M. nasutus* pseudoreference | 82.2 | 4.1 | 0.3 |
| RK23 | $RSB_7$ STE | carpel | STEcp | 2 | 16 | 160 | 16,482,294 | 16,429,628 | 0.32 | *M. nasutus* pseudoreference | 73.6 | 3.9 | 0.3 |
| RK24 | $RSB_7$ STE | carpel | STEcp | 3 | 16 | 229 | 14,508,347 | 14,459,916 | 0.33 | *M. nasutus* pseudoreference | 73.8 | 4.0 | 0.3 |

**Table S2. Hybrid gene expression categories.** Gene expression was categorized across carpels and stamens of fertile (FER) and sterile (STE) hybrid siblings from a seventh-generation SF5-IM62 backcross population, *r* ecurrent *s* election with *b* ackcrossing (RSB7) based on three pairwise comparisons: (i) *M. nasutus* SF5 versus *M. guttatus* IM62 parents, (ii) hybrid versus SF5, and (iii) hybrid versus IM62. S/NS = difference in transcript abundance between samples is significant (S, -2<log2 fold-change>2, FDR<=0.05) or non-significant (NS) within each comparison. Hybrid = FER or STE RSB7 siblings. UP/DOWN = transcript abundance is higher (UP) or lower (DOWN) in SF5 compared to IM62.

| Category | SF5 vs IM62 | Hybrid vs SF5 | Hybrid vs IM62 | Color |
|---|---|---|---|---|
| Conserved | NS | NS | NS | grey |
| | | NS | S | |
| | | S | NS | |
| IM62-like conserved | NS | S | NS | yellow |
| IM62-like divergent | S | S | NS | orange |
| Intermediate | S | NS | NS | green |
| | | S DOWN | S UP | |
| | | S UP | S DOWN | |
| SF5-like conserved | NS | NS | S | light blue |
| SF5-like divergent | S | NS | S | dark blue |
| Transgressive conserved | NS | S UP | S UP | light pink |
| | | S DOWN | S DOWN | |
| Transgressive divergent | S | S UP | S UP | dark pink |
| | | S DOWN | S DOWN | |

**Table S3. GO term enrichment among parental stamen-biased genes and STE stamen downregulated genes.** Shown are the top 10 biological processe-related GO terms for two categories of genes: (i) downregulated in stamens of the sterile (STE) genotypes from the RSB7 introgression population and (i) stamen-biased in the parents, *Mimulus nasutus* SF5 and *Mimulus guttatus* IM62. Of the 2192 downregulated STE stamen genes, 1405 had GO term assignments. Of the 2038 parental stamen-biased genes, 1585 had GO term assignment.

| GO term ID | GO Term | Annotated | Count | Expected | p-value | q-value |
|---|---|---|---|---|---|---|
| *STE stamen downregulated genes* | | | | | | |
| GO:0005975 | carbohydrate metabolic process | 842 | 129 | 63.4 | 9.60E-16 | 4.59E-12 |
| GO:0009860 | pollen tube growth | 71 | 29 | 5.4 | 5.50E-15 | 1.31E-11 |
| GO:0009932 | cell tip growth | 82 | 31 | 6.2 | 8.20E-15 | 1.31E-11 |
| GO:0055085 | transmembrane transport | 732 | 112 | 55.2 | 1.10E-13 | 1.31E-10 |
| GO:0044703 | multi-organism reproductive process | 220 | 50 | 16.6 | 8.00E-13 | 7.65E-10 |
| GO:0048588 | developmental cell growth | 98 | 31 | 7.4 | 2.20E-12 | 1.75E-09 |
| GO:0048868 | pollen tube development | 101 | 31 | 7.6 | 5.30E-12 | 3.62E-09 |
| GO:0045229 | external encapsulating structure organization | 252 | 52 | 19.0 | 1.40E-11 | 8.36E-09 |
| GO:0071555 | cell wall organization | 232 | 49 | 17.5 | 2.30E-11 | 1.22E-08 |
| GO:0000904 | cell morphogenesis involved in differentiation | 148 | 36 | 11.2 | 1.80E-10 | 8.60E-08 |
| *Parental stamen-biased genes* | | | | | | |
| GO term ID | GO Term | Annotated | Count | Expected | p-value | q-value |
| GO:0055085 | transmembrane transport | 732 | 128 | 65.0 | 1.40E-14 | 6.69E-11 |
| GO:0051234 | establishment of localization | 1672 | 233 | 148.4 | 6.00E-14 | 1.27E-10 |
| GO:0051179 | localization | 1723 | 238 | 152.9 | 8.00E-14 | 1.27E-10 |
| GO:0006810 | transport | 1654 | 230 | 146.8 | 1.20E-13 | 1.43E-10 |
| GO:0009860 | pollen tube growth | 71 | 29 | 6.3 | 3.80E-13 | 3.63E-10 |
| GO:0005975 | carbohydrate metabolic process | 842 | 137 | 74.7 | 4.80E-13 | 3.82E-10 |
| GO:0009932 | cell tip growth | 82 | 31 | 7.3 | 7.00E-13 | 4.78E-10 |
| GO:0044765 | single-organism transport | 1274 | 184 | 113.1 | 2.40E-12 | 1.43E-09 |
| GO:1902578 | single-organism localization | 1303 | 186 | 115.6 | 5.20E-12 | 2.76E-09 |
| GO:0048868 | pollen tube development | 101 | 32 | 9.0 | 7.00E-11 | 3.35E-08 |
| GO:0015672 | monovalent inorganic cation transport | 175 | 44 | 15.5 | 1.20E-10 | 5.21E-08 |

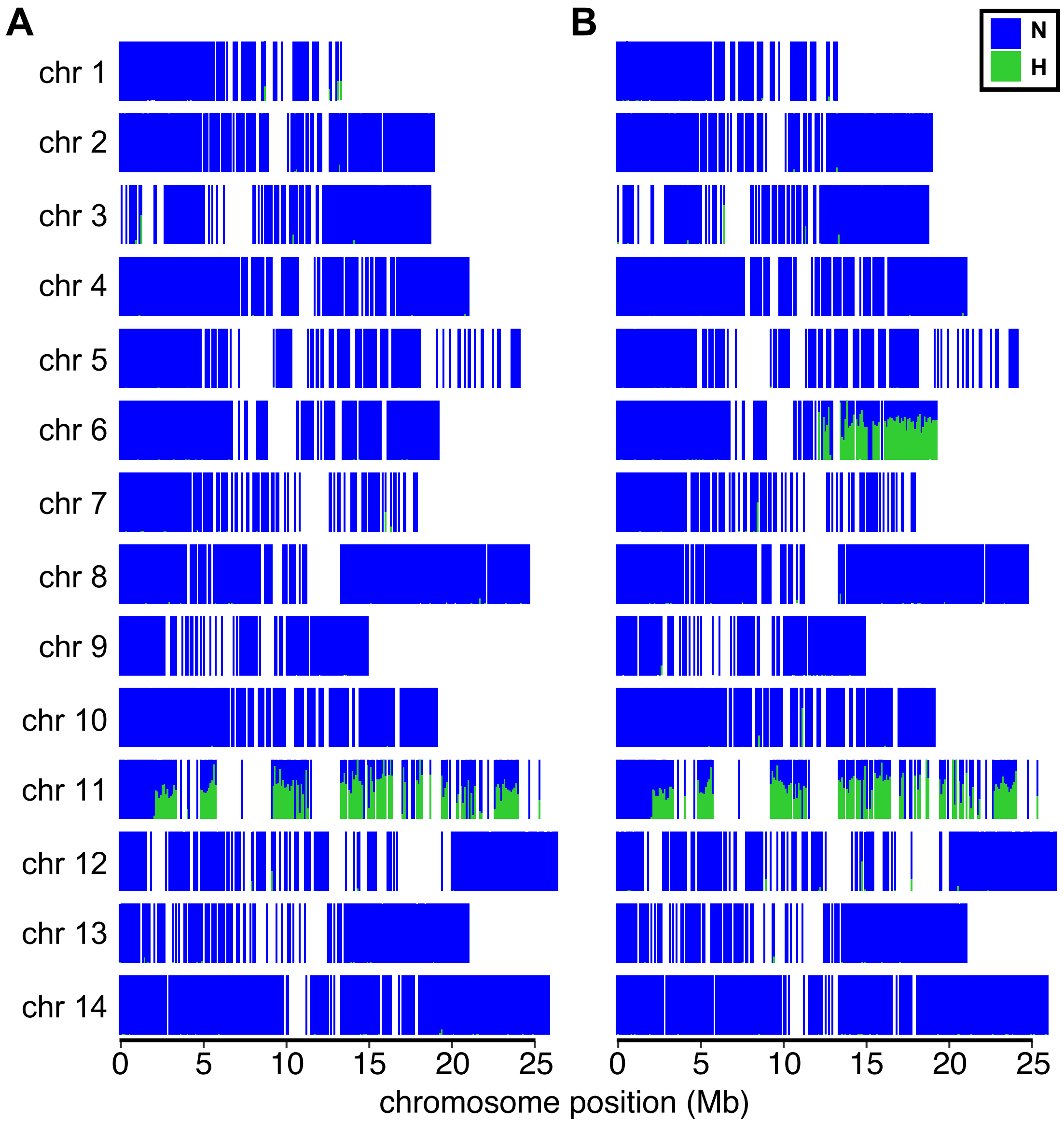


**Figure S1. Genome-wide parental ancestry of (A) fertile (FER) and (B) sterile (STE) RSB$_7$ siblings**. Plots show the proportion of heterozygous *M. guttatusIM62-M. nasutus* SF5 (H, green) and homozygous SF5 (N, blue) ancestry in 50Kb bins across the 14 chromosomes. The length of each chromosome represents physical size according to the *M. guttatus* v2.0 assembly. Genotype calls are based on over 250,000 biallelic single nucleotide polymorphisms (SNPs). Chromosomal regions without reliable SNP calls are represented by white space. FER and STE genotypes carry a heterozygous introgression along 23 Mb of chromosome 11 associated with a meiotic drive element (*D*). STE genotypes carry an additional 7 Mb heterozygous introgression around *hms1* on chromosome 6.

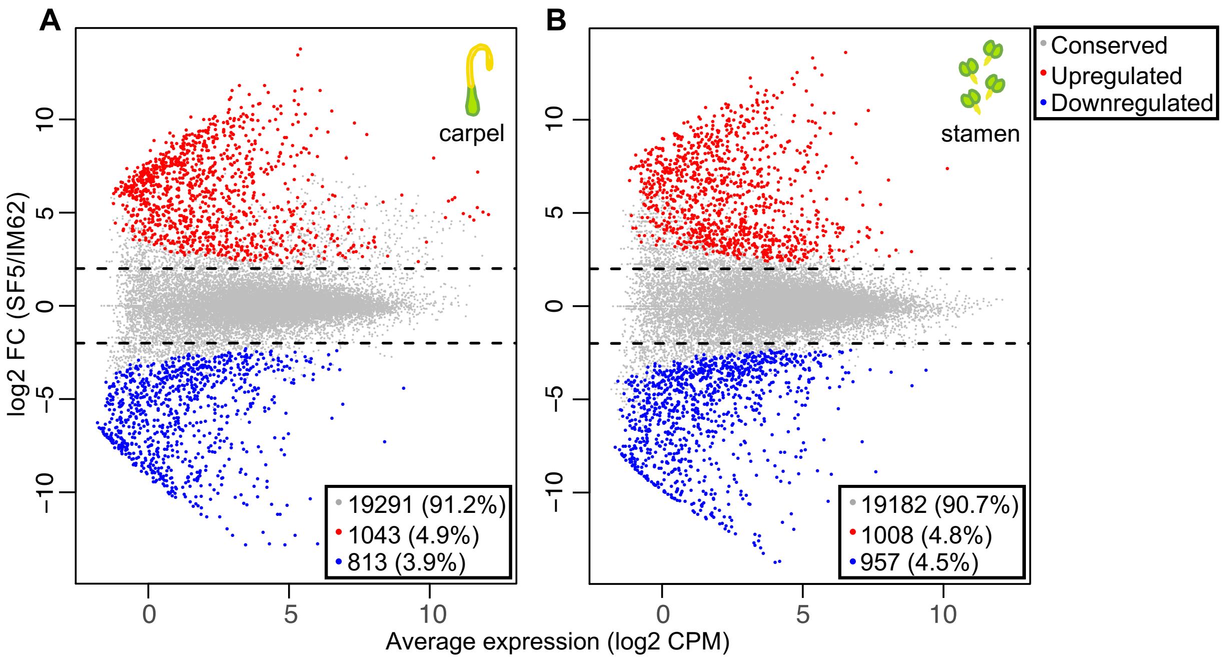


**Figure S2. Interspecific regulatory divergence between SF5 and IM62.** Plot shows relative transcript abundance (log2 fold-change (FC)) between the parents, *Mimulus nasutus* SF5 and *M. guttatus* IM62, versus average transcript abundance (log2 counts-per-million (CPM)) in (A) carpels and (B) stamens. Colors indicate whether transcript abundance is conserved (grey) or significantly (FDR≤0.05) downregulated (blue) or upregulated (red) in SF5 versus IM62. Dashed lines at 2 and -2 mark the significance thresholds for upregulation and downregulation, respectively.

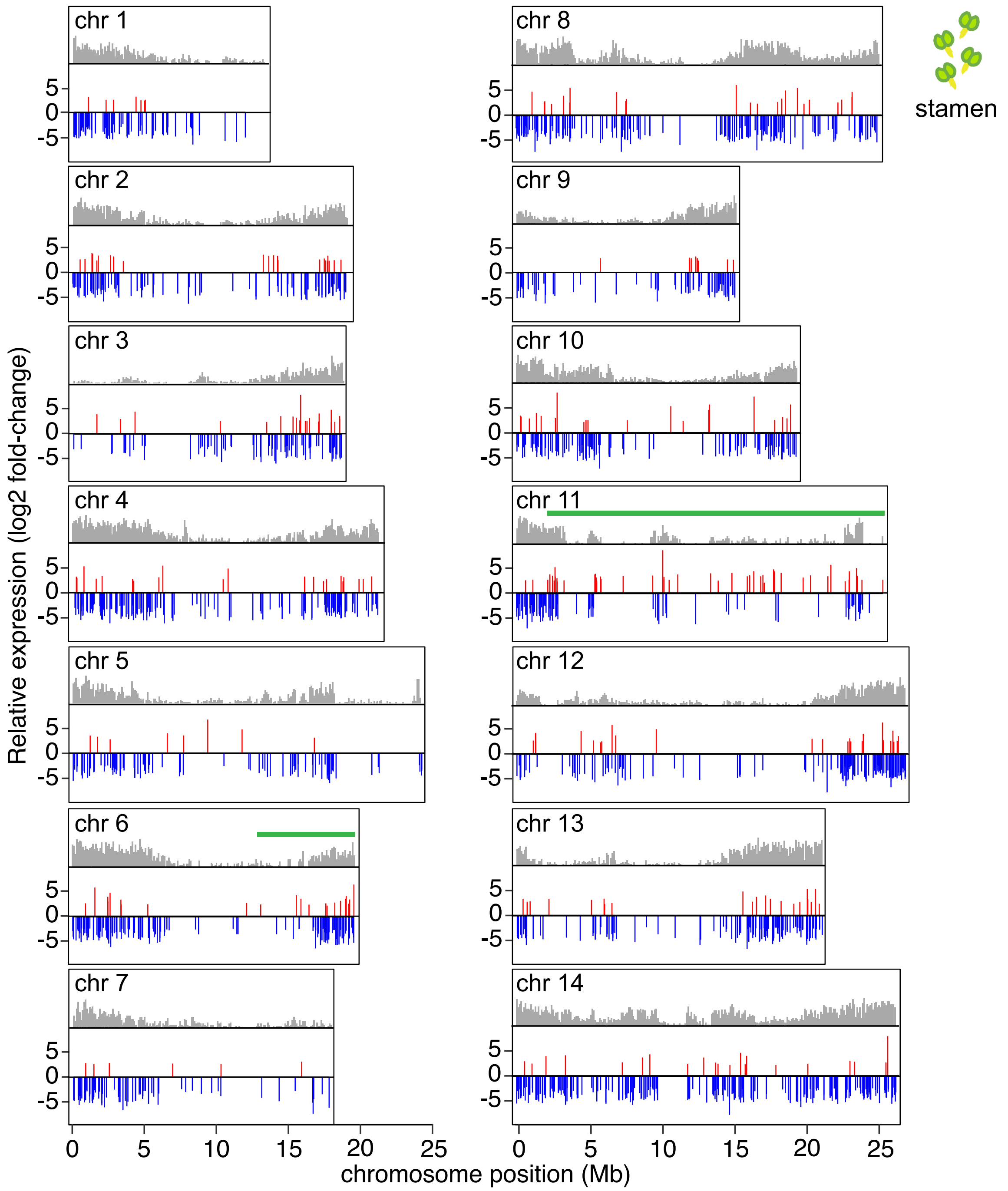


**Figure S3. Genome-wide distribution of differentially expressed genes (DEGs) in STE stamens.** Plot shows relative transcript abundance (log2 fold-change (FC)) across the 14 *Mimulus* chromosomes for the 2192 DEGs (-2<log2 FC>2, FDR≤0.05) in STE stamens compared to FER or SF5 stamens. To identify DEGs in the heterozygous introgressions and homozygous background regions, transcript abundance was compared to the SF5-IM62 mid-parent value or SF5 parent, respectively. Grey histograms represent relative density of expressed genes in 50Kb bins across the 14 chromosomes. Green bars demarcate the heterozygous introgressions on chromosomes 6 and 11.

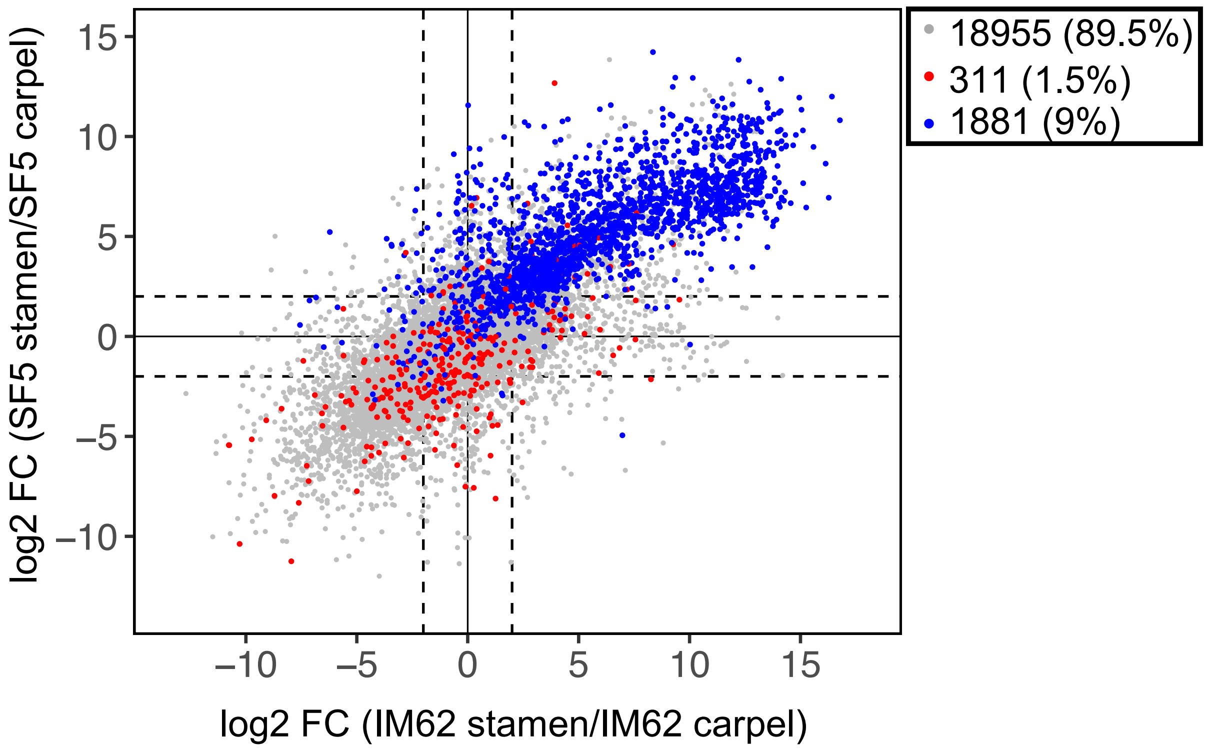


**Figure S4. Overrepresentation of stamen-biased genes among differentially expressed genes (DEGs) in STE stamen.** Scatterplot shows the pattern of parental tissue-biased expression (log2 fold-change (FC)) for genes that were significantly upregulated (log2 FC>2, FDR≤0.05; red), downregulated (-2<log2 FC, FDR≤0.05; blue) or conserved (grey) in STE stamens compared to FER or SF5 stamens. Of the 2192 STE stamen DEGs, 1372 and 55 were stamen- and carpel-biased, respectively, in the parents.

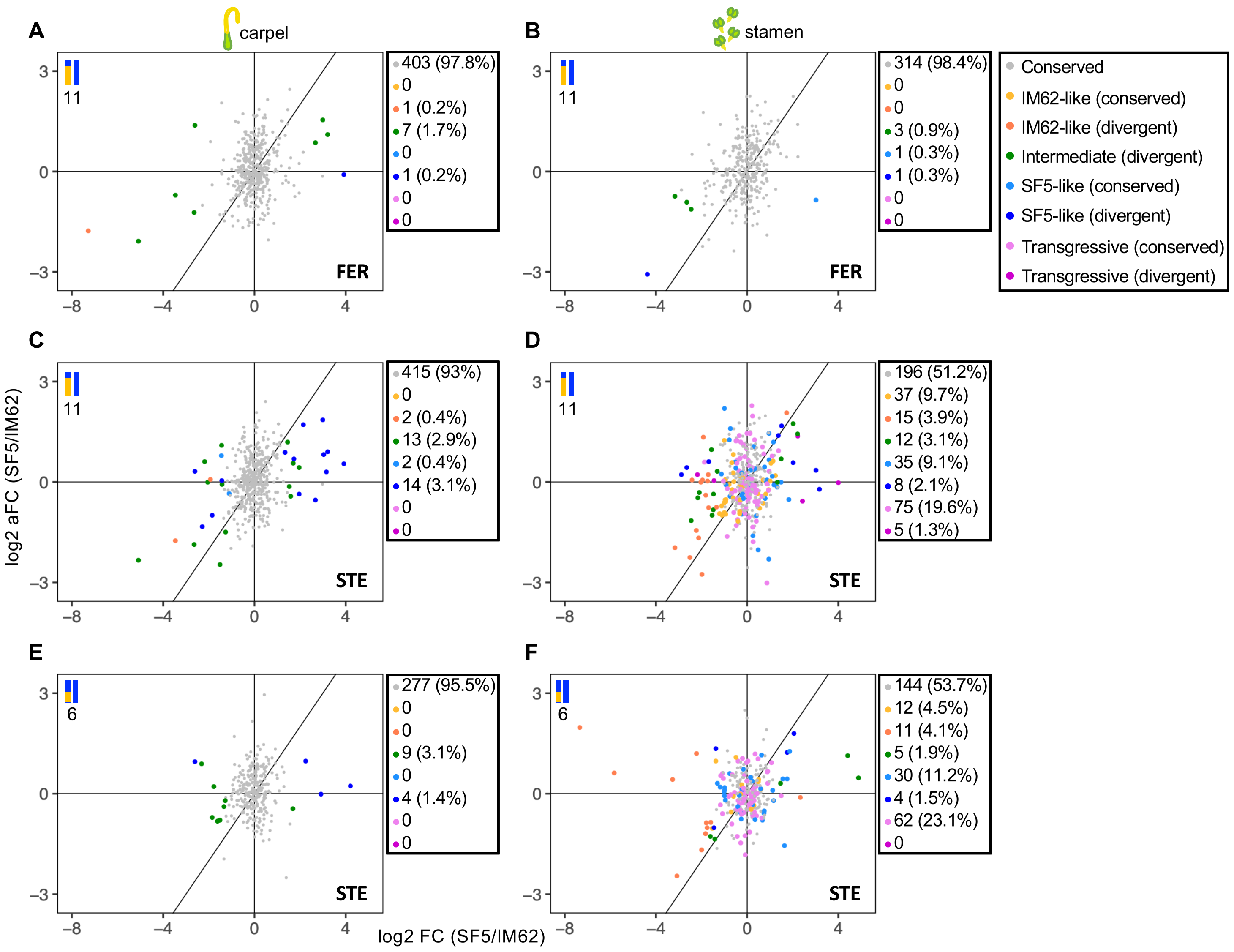


**Figure S5. Pattern of *cis*- and *trans*-regulatory differences in FER and STE introgression regions.** Plots show relative transcript abundance (log2 fold-change (FC)) between the parents on the x-axis and relative allelic transcript abundance (log2 allelic FC (aFC) within heterozygous introgression regions on the y-axis. This figure is identical to Figure 7 except that here the genes are colored by expression category (see Table S2 for description).
